# Senescence of stromal cells contributes to endometrium dysfunction and embryo implantation failure

**DOI:** 10.1101/2021.07.19.452880

**Authors:** Pavel I. Deryabin, Julia S. Ivanova, Aleksandra V. Borodkina

## Abstract

Successful implantation requires two-sided interaction between genetically normal embryo and properly prepared endometrium. At the cellular level the latter means hormone-regulated differentiation of endometrial stromal cells (ESCs) into decidual ones that create appropriate microenvironment for invading embryo. Impaired decidualization is proved to mediate implantation failures. Here we elicited ESCs’ senescence as the cause for disturbed decidualization of endometrial stroma and impaired blastocyst implantation. Ability to decidualize and to accept modeled blastocysts inversely correlated with senescence in patients’ ESCs lines. Reduced hormonal responsiveness of senescent ESCs led to inappropriate decidualization dynamics resulting in altered receptivity, disturbed ligand-receptor interaction with trophoblasts and modified architecture of extracellular matrix what hindered blastocysts’ invasion. Furthermore, senescent ESCs caused ‘bystander’ quenching of decidual reaction in adjacent cells reinforcing dysfunction of stromal compartment. Implementation of senomorphics reducing senescence phenotype diminished adverse effects of senescent ESCs on decidualization and implantation using both in vitro models and patients’ lines, what suggests a promising strategy to increase in vitro fertilization efficacy.

## Introduction

The more data on cell senescence appear, the more difficult it becomes to find an unambiguous definition for this phenomenon. From the very beginning senescence could be simply defined as an irreversible proliferation loss of cells with uncapped telomere ends (Hayflick, 1965; Campisi, 1996). In contrast to apoptotic cells, senescent cells preserve viability and metabolic activity though harboring severe intracellular alterations (Campisi, 1996). Depending on the inducer’s nature the panel of senescence forms was substantially extended. Today stress-induced, oncogene-induced, tumor suppressor-induced, therapy-induced forms of senescence are described along with the replicative one (Campisi, 1996; Serrano et al, 1997; Toussaint et al, 2000; Roninson et al, 2001; Peeper, 2010; Demaria et al, 2017). Furthermore, senescence was initially considered as a response inherent only to proliferating cells. However, more recent findings elicit that some post-mitotic cells are also capable for senescence (von Zglinicki et al, 2020; Sapieha & Mallette, 2018). Bearing all these data in mind, the term ‘senescence’ became far more complex compared to its original version.

Simultaneously with the expansion of the term ‘senescence’ its functional outcomes were also revisited. Firstly, senescence was considered as an intrinsic anti-tumor barrier that prevented proliferation of cells bearing damages (Campisi et al, 1996). Next, it was shown that senescence-associated secretory phenotype (SASP) produced by senescent cells can facilitate tumor development in the preneoplastic surrounding, suggesting its positive role in cancer progression (Coppé et al, 2010). Further, accumulation of senescent cells within tissues was proved to mediate their dysfunction and progression of various pathologies (Franceschi & Campisi, 2014; Borghesan et al, 2020; Karin & Alon, 2021). Contrarily, short-term presence of senescent cells appeared to be essential for optimal wound healing, tissue repair and regeneration, as well as during embryonic development (Demaria et al, 2014; Walters & Yun, 2020; Storer et al, 2013). Today the role of cellular senescence is extensively studied with regard to various pathologies (He & Sharpless, 2017; Borghesan et al, 2020; Karin & Alon, 2021). Promising results in treating diseases of different etiologies have been already obtained based on the senolytics or senomorphics applications, leading either to targeted killing of senescent cells or to reduction of SASP secretion (Song et al, 2020). However, from this point of view, endometrium and female infertility still remain poorly studied, probably, due to several obstacles, which will be discussed below.

The obvious intricacy considering endometrium investigation is the complex dynamic nature of this tissue regulated by the multilevel hormonal networks (Deryabin et al, 2020; Critchley et al, 2020). Endometrium is an inner lining of the uterus that consists of two layers – basalis and functionalis. The crucial cellular components of both layers are endometrial stromal cells (ESCs). Each menstrual cycle starts with the proliferative/follicular phase during which ESCs from basalis actively proliferate forming new functional layer. Simultaneously, the dominant follicle matures in the ovary until the ovulation that occurs in the middle of the menstrual cycle. Ovulation results in the release of an egg from the ovary into fallopian tube. Subsequently, corpus luteum producing progesterone is formed at the site of the dominant follicle. This marks the onset of the second secretory/luteal phase of the menstrual cycle. During this phase functional layer of the endometrium undergoes crucial transformation governed mainly by progesterone synthesized by the corpus luteum. Such endometrial transformation is termed decidualization and at the cellular level presents tissue-specific differentiation of ESCs into decidual cells (Park et al, 2016; Yoshie et al, 2015; Okada et al, 2018). Proper decidualization mediates so-called ‘window of implantation’ (WOI) – short time-period when endometrial tissue becomes receptive and enables embryo implantation. Decidual cells are essential for trophoblast invasion and growth, prevention of maternal immunological rejection, promotion of angiogenesis, and thus for the establishment of pregnancy (Gellersen & Brosens, 2014; Okada et al, 2018; Deryabin et al, 2020). In the absence of fertilization decidualized functional layer of the endometrium sheds and new proliferative phase begins.

Despite of the obvious importance of proper endometrial functioning for embryo implantation, during rather long time period implantation failures were rarely associated with endometrial factor. In this context, the main focus was and still is on the ovarian reserve and embryo quality. Largely due to the introduction of preimplantation genetic testing for aneuploidy (PGT-A) during in vitro fertilization (IVF), the impact of endometrium into the success of implantation became more evident (Tomari et al, 2020). Today about one-third of implantation failures is regarded to be mediated by inadequate endometrial receptivity (Altmäe et al, 2017; Tomari et al, 2020).

Another important aspect regarding endometrial studying that also cannot be ignored is the inappropriateness of the common animal models. Together with humans, only higher primates, some species of bats, and the elephant shrew have menstruation, while most other mammals including mice and rats have estrous cycle instead of the menstrual one (Emera et al, 2012). Besides for bleeding, there are other crucial differences between these two types of cycling, among which ESCs decidualization that begins only after embryo implantation in estrous cycle, while precedes implantation during menstrual cycle.

Together these complexities might partially explain the minor interest in studying the role of cellular senescence in endometrial functioning and female fertility. Nevertheless, recently, the data regarding the existence of the relationship between ESCs decidualization and senescence began to appear (Lucas et al, 2016a; Lucas et al, 2016b; Brighton et al, 2017; Cha & Aronoff, 2017; Marquez et al, 2017; Durairaj et al, 2017; Tomari et al, 2020). In particular, it was shown that altered ESCs secretome, in many ways similar to SASP, preceded implantation failure (Durairaj et al, 2017). Based on the DNA methylation analysis senescent ESCs were suggested to be partially responsible for decreased endometrial plasticity and thus might mediate recurrent pregnancy losses (Lucas et al, 2016a). Furthermore, it was speculated that implantation failure might be associated with enhanced level of ESCs senescence during the proliferative phase of the menstrual cycle (Tomari et al, 2020). Despite of the certain prerequisites that suggest the role of senescence in endometrium, today there is a lack of comprehensive understanding of how ESCs senescence might influence endometrial tissue functioning. Therefore, the aim of the present study was to investigate the contribution of ESCs senescence to decidual reaction and embryo implantation.

## Results

### 1. Decidual reaction of the primary ESCs lines from IVF patients is inversely related to the degree of senescence

To reveal possible biological role of ESCs senescence in endometrium functioning, we compared several primary ESCs lines by the level of senescence, on the one hand, and by the ability to differentiate into decidual cells, on the other. ESCs lines were obtained from patients planning to undergo IVF. Of note, ESCs isolation procedure was unified with regard to the phase of the menstrual cycle and the absence of any endometrial complications.

In order to rank ESCs lines by the level of senescence, we estimated the following senescence-related parameters: cell size, lipofuscin accumulation, p21 expression and SA-β-Gal activity. Interestingly, primary ESCs lines varied significantly by the level of all the tested senescence markers, suggesting different degree of senescence (Fig EV1A–D).

Since decidualization is the main physiological role of ESCs that mediates hormonal responsiveness of endometrial tissue, we next assessed the ability of the same patients’ lines to decidualize. To induce decidual differentiation, we used the classical cocktail containing cAMP, β-estradiol (E2), and synthetic progesterone analog medroxyprogesterone 17-acetate (MPA) (Deryabin et al, 2021). To compare decidual response between the tested ESCs lines, we estimated expression of the key transcription factor *FOXO1* and decidual marker genes – *PRL*, *IGFBP1* and *CLU* (Fig EV1E–H). Additionally, we applied genetic tool that reflects functioning of the core decidual network by the fluorescence intensity of the reporter protein, which was designed by us and described in detail in our previous study (Deryabin et al, 2021). According to our data, the intensity of the decidual reaction varied greatly between ESCs lines (Fig EV1I).

The fact that cell lines obtained from different patients demonstrated different degree of senescence markers and decidualization may seem not so surprising itself. Far more important is that we were able to reveal negative correlation between the basal level of senescence and decidualization ability, i.e. the more pronounced senescence markers were, the worse differentiation potential of such cells was (Fig 1). The extreme variants are line 2304 characterized by the maximal elevation of all the tested senescence parameters and the significantly reduced decidual response, and line 1410 with the most pronounced decidual reaction and minimal if any senescence signs. Together these data evidence in favor of the negative role of ESCs senescence in endometrium functioning and provide basis for further detailed investigation of the molecular mechanisms of this influence.

**Figure 1.**
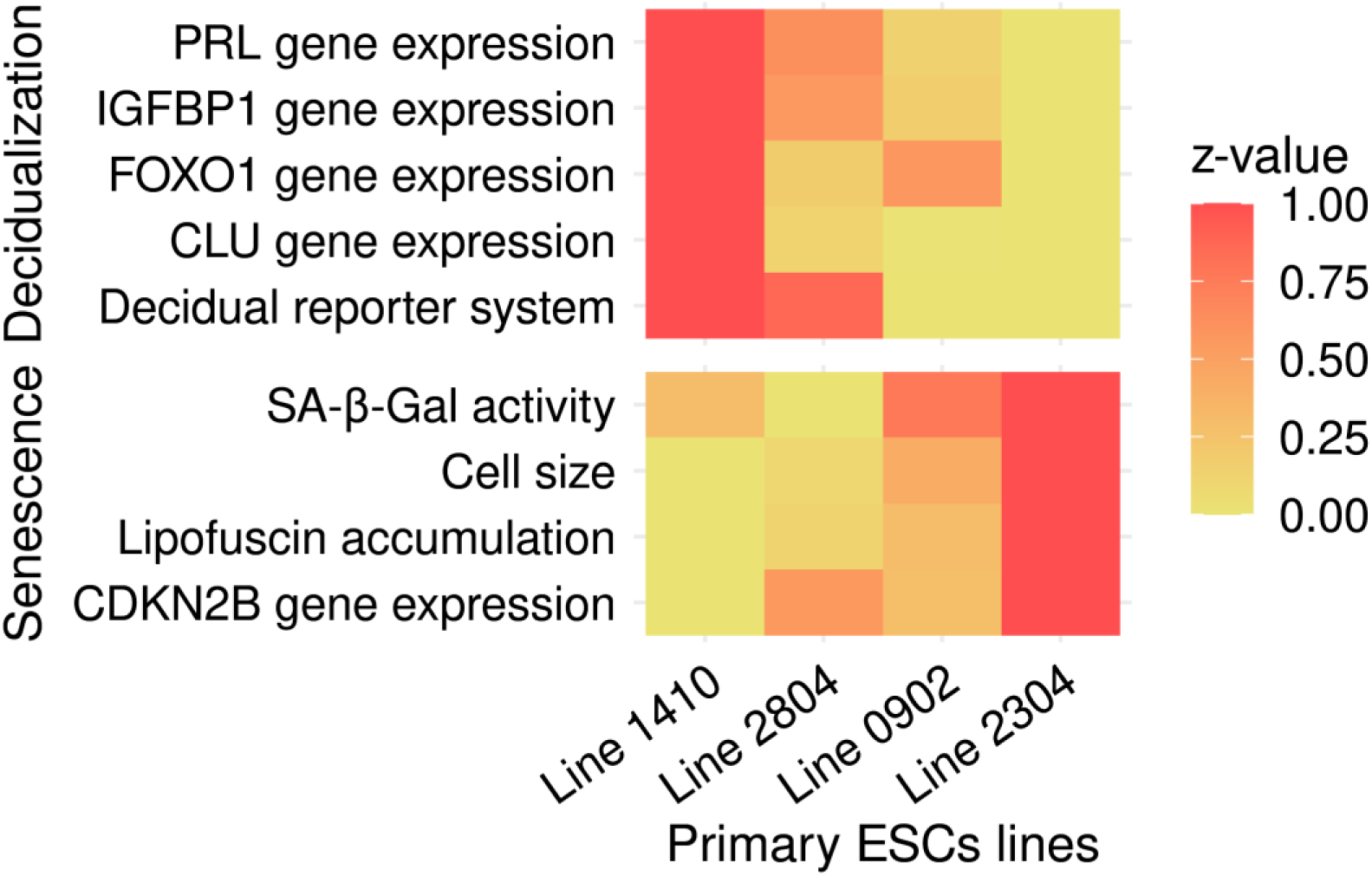
Correlation between the basal level of senescence and decidualization ability of the primary ESCs lines. Data information: Data are presented as max-min normalized values; the original values for each parameter are provided Fig EV1.

### 2. Senescent ESCs have impaired decidual response upon hormonal stimulation

The initial suggestion of how ESCs senescence may influence endometrial tissue functioning was based on the assumption that senescent cells might have impaired differentiation properties. Thus, we assessed the ability of senescent ESCs to decidualize in response to hormonal supplementation. To this end, we applied in vitro model of oxidative stress-induced senescence of ESCs described in detail in our previous studies and compared decidual reactions of young and senescent ESCs treated with the hormonal cocktail (Burova et al, 2013; Borodkina et al, 2014; Griukova et al, 2019). Of note, ESCs treated with sublethal oxidative stress displayed all the common features of senescent cells, including irreversible cell cycle block, complete proliferation loss, activation of p53/p21 or p16/Rb pathways, cellular hypertrophy, increased lipofuscin accumulation, persistent DNA damage, activity of senescence-associated β-galactosidase (SA-β-Gal), impaired mitochondrial functioning, increased intracellular reactive oxygen species (ROS) levels, and senescence-associated secretory phenotype (SASP) (Borodkina et al, 2014; Griukova et al, 2019). As shown in Fig. 2a, b, young ESCs switched morphology from fibroblast-like to polygonal epithelial-like what is typical for decidualization, while morphology of senescent cells remained almost unchanged (Fig 2A and B). In line with this observation, senescent cells displayed less pronounced alterations in the expression dynamics of E-cadherin and vimentin, suggesting impaired mesenchymal-to-epithelial transition (Fig 2E). Also, we revealed significantly reduced expression of the core decidual transcription factor *FOXO1* along with the decreased mRNA levels of the decidual marker genes – *PRL* and *IGFBP1* (Fig 2C). Prolactin is secreted by the decidual cells and contribute to trophoblast growth and angiogenesis (Deryabin et al, 2020). As expected, senescent ESCs secreted lower amounts of prolactin upon decidualization compared to the young cells (Fig 2D). The proper hormonal response of ESCs is governed primarily by the two steroid hormone receptors – progesterone (PR) and estrogen (ER), whose expression increased substantially during decidualization of young ESCs (Fig 2E). However, expression of both receptors was slightly above the basal level in senescent cells upon hormonal supplementation (Fig 2E). Upon binding to progesterone PR translocates into the nucleus, where it recognizes and binds to specific DNA sequences termed progesterone responsive elements, and thus directly regulates the expression of a large number of decidual genes (Deryabin et al, 2021). Indeed, immunofluorescent staining with PR antibodies clearly indicates PR translocation in young decidualized ESCs, while in senescent cells PR distribution in undifferentiated and differentiated states was similar (Fig 2F). It should be specifically highlighted that the disturbed decidualization of senescent ESCs was additionally verified using another senescence model – the replicative one. The main features of both types of senescent ESCs were described in our previous studies (Deryabin & Borodkina, in press; Deryabin et al, preprint). Similar to the results presented above, replicatively senescent ESCs demonstrated impairments of the proper hormonal responsiveness (Fig EV2). The data obtained clearly indicate disturbed ability of senescent ESCs to decidualize in response to hormonal stimulation.

**Figure 2.**
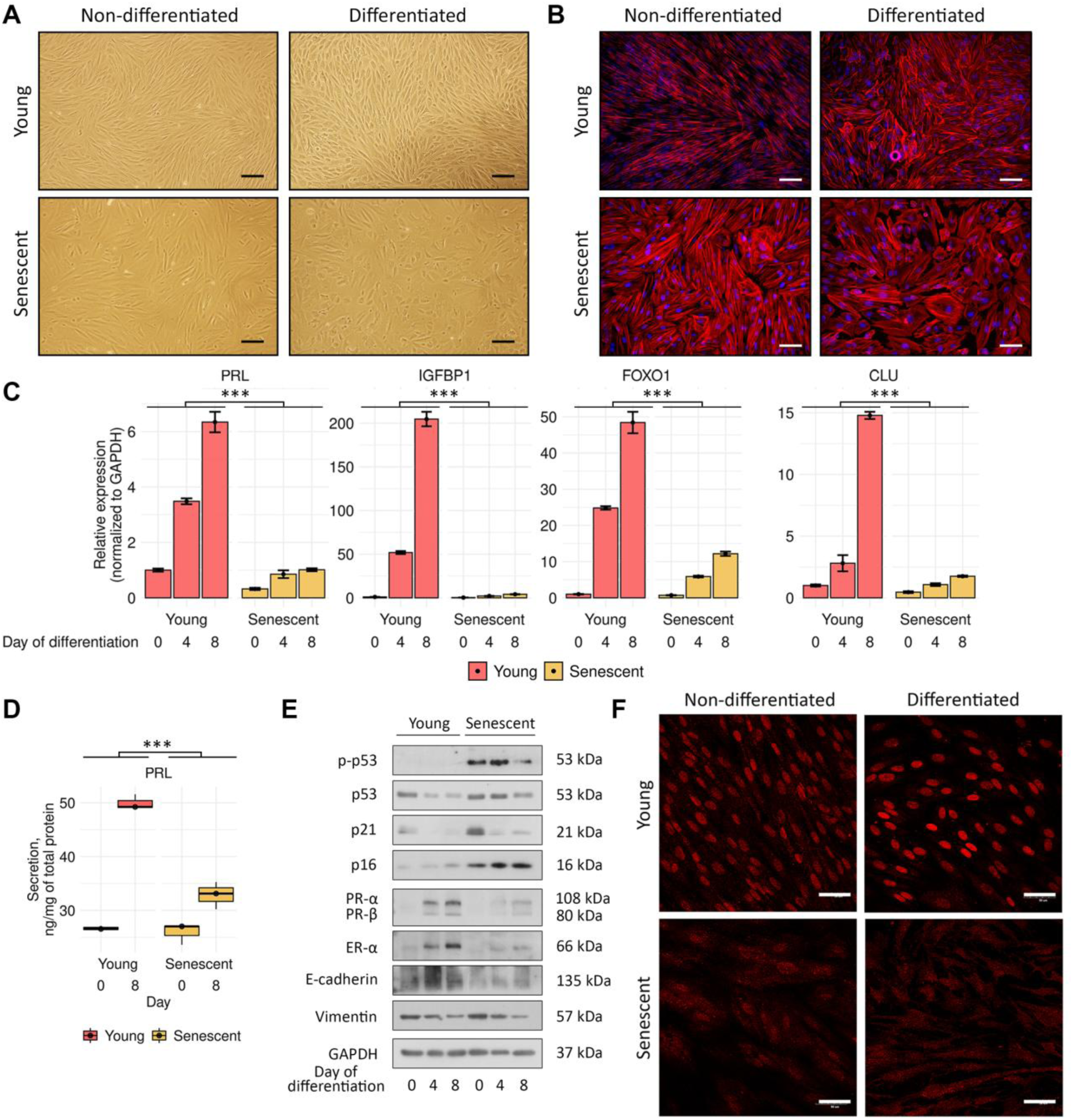
Impaired decidual differentiation of senescent ESCs. A, B Disturbed morphology of senescent ESCs upon decidualization observed in bright-field (A) and by F-actin cytoskeleton visualization (B). Scale bars: 100 μm. C Quantitative RT-PCR detection of *PRL*, *IGFBP1*, *FOXO1*, *CLU* expression levels in young and senescent ESCs (n = 3 for each sample). D Prolactin secretion assessment in young and senescent ESCs (n = 3 for each sample). E Expression and phosphorylation levels of the proteins related to senescence (pp53, p21, p16), to decidualization (PR and ER) and to mesenchymal-to-epithelial transition (E-cadherin, vimentin) in undifferentiated and decidualized young and senescent ESCs. F Immunofluorescent staining with PR antibodies. Scale bars: 50 μm. Data information: In (C), data are presented as mean ± SD. In (D), data are presented as median ± IQR. ***P<0.005 (two-sided ANOVA).

### 3. Time-course RNA-seq analysis revealed altered decidualization dynamics in senescent ESCs

In order to detail molecular differences that occur during decidualization of young and senescent ESCs, we performed RNA-seq analysis. The starting point for the analysis was day 0 (undifferentiated cells); the following time points were 4 and 8 days of differentiation. Principal component analysis clearly demonstrated different expression patterns for young and senescent cells (PC2) both before the induction of decidualization (PC1) and in each time point during differentiation (Fig 3A). By applying LRT-test we revealed 2963 (FDR<0.01) differentially expressed genes (DEGs) between young and senescent cells in course of decidualization (Table 1). Based on the levels of genes expression and the direction of their alterations 2932 of the identified DEGs were further clustered into 21 groups that included not less than 19 genes (Fig 3B). The obtained clusters provided clear illustration of the distinct decidualization dynamics between young and senescent ESCs. The following functional annotation of these clusters in Gene Ontology (GO) terms for Biological Processes (BP) uncovered altered responsiveness to hormones and impairments in steroid biosynthetic processes, disturbed communication with immune, epithelial and endothelial cells, improper angiogenesis and disorganized extracellular matrix during decidualization of senescent ESCs (Fig 3C). Correct progression of these processes is crucial for proper functioning of endometrial tissue. The multifaceted analysis of each cluster as well as of unclustered genes is provided in Appendix FigS1–22 and in Table1. Summarizing the data obtained, we can conclude that decidualization progression in senescent ESCs differs significantly from that in the young ones, what should certainly affect the implantation process.

**Figure 3.**
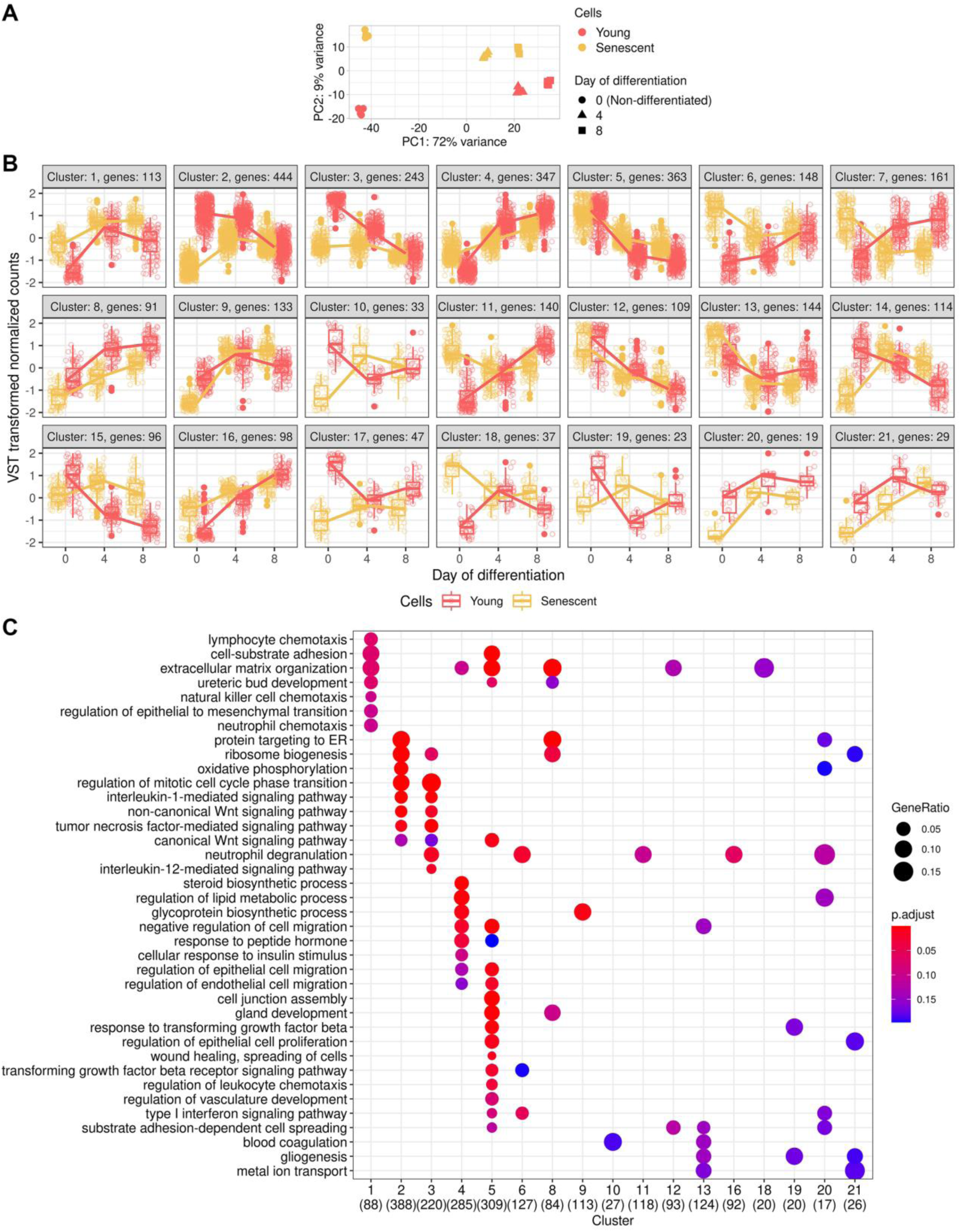
Comparative analysis of the decidualization dynamics between young and senescent ESCs. A PCA plot displaying global gene expression profiles in young and senescent ESCs during decidualization progression (n = 4 for each condition). B Clusterization of the identified DEGs based on the levels of genes expression and the direction of their alterations. Each cluster included not less than 19 genes. C Functional annotation of the identified clusters in GO:BP terms. Data information: In (B), expression values are presented as VST transformed normalized counts.

**Table 1.**
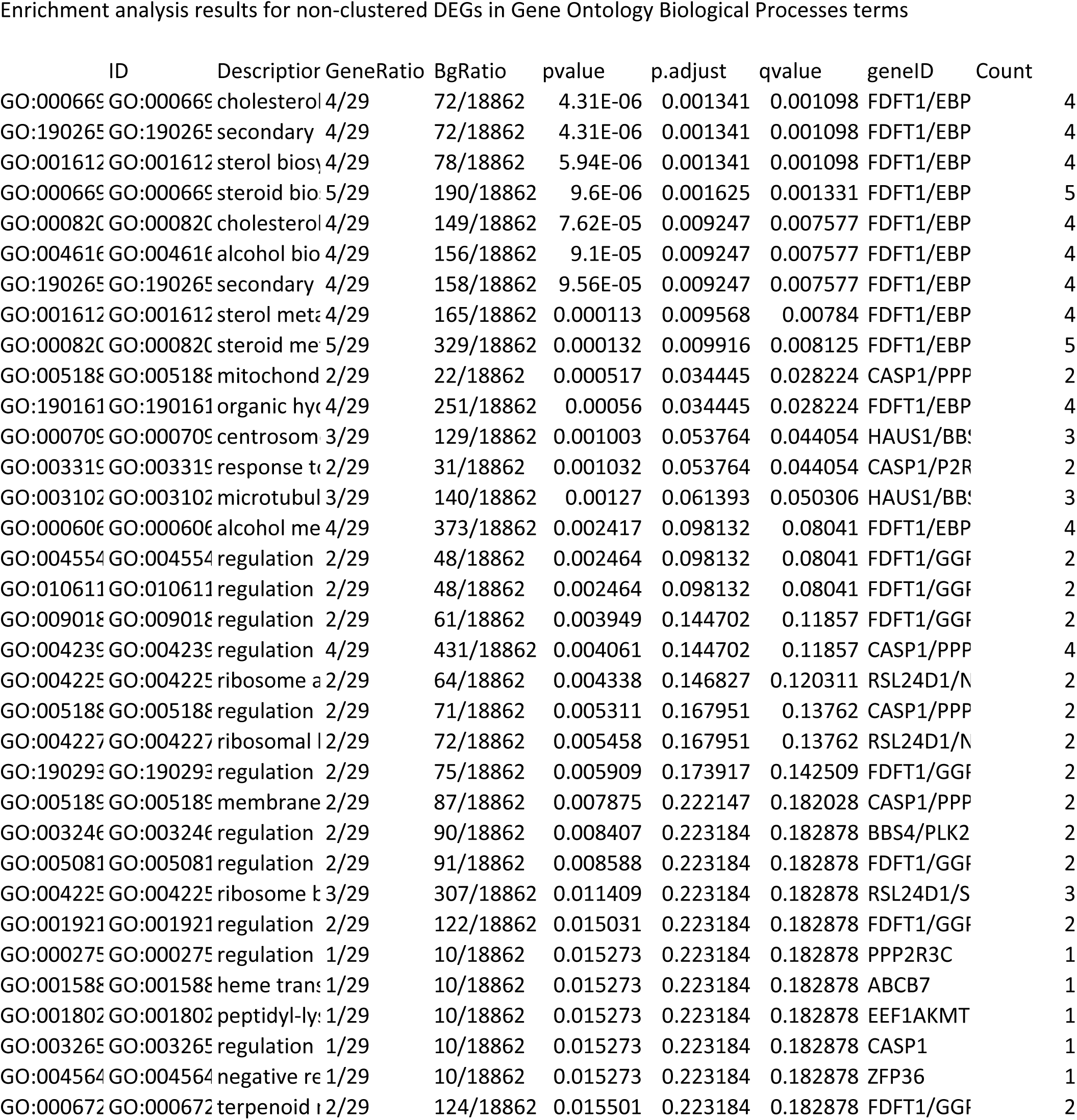
Extended results of bioinformatic analysis of RNA-Seq data. List1 Results of differential gene expression estimation among all six groups in the dataset (LRT-test). List2 Differentially expressed genes (LRT-test p.adj < 0.01) from List1 subjected to clusters. List3–24 Enrichment analysis results for Clusters form 1 to 21 and for non-clustered DEGs in Gene Ontology Biological Processes terms.

### 4. Inability of senescent ESCs to decidualize properly upon hormonal stimuli mediates impaired blastocysts invasion

The correct response of ESCs to the increasing levels of steroid hormones during the second phase of the menstrual cycle resulting in decidual transformation of endometrial tissue marks WOI (Deryabin et al, 2020). During this limited time period endometrium becomes receptive and enables further implantation. In the recent study the ‘meta-signature’ of human endometrial receptivity was unraveled (Altmäe et al, 2017). The authors identified 57 mRNA genes as putative receptivity markers. Using our RNA-seq data we analyzed the enrichment of this ‘receptivity subset’ among DEGs during decidualization of young and senescent ESCs. As shown in Fig 4A, senescent ESCs were characterized by disturbed expression of the most receptivity markers (FDR 1.58e-06), what should negatively affect the overall receptivity of the endometrial tissue, thus creating unfavorable microenvironment for embryo implantation.

**Figure 4.**
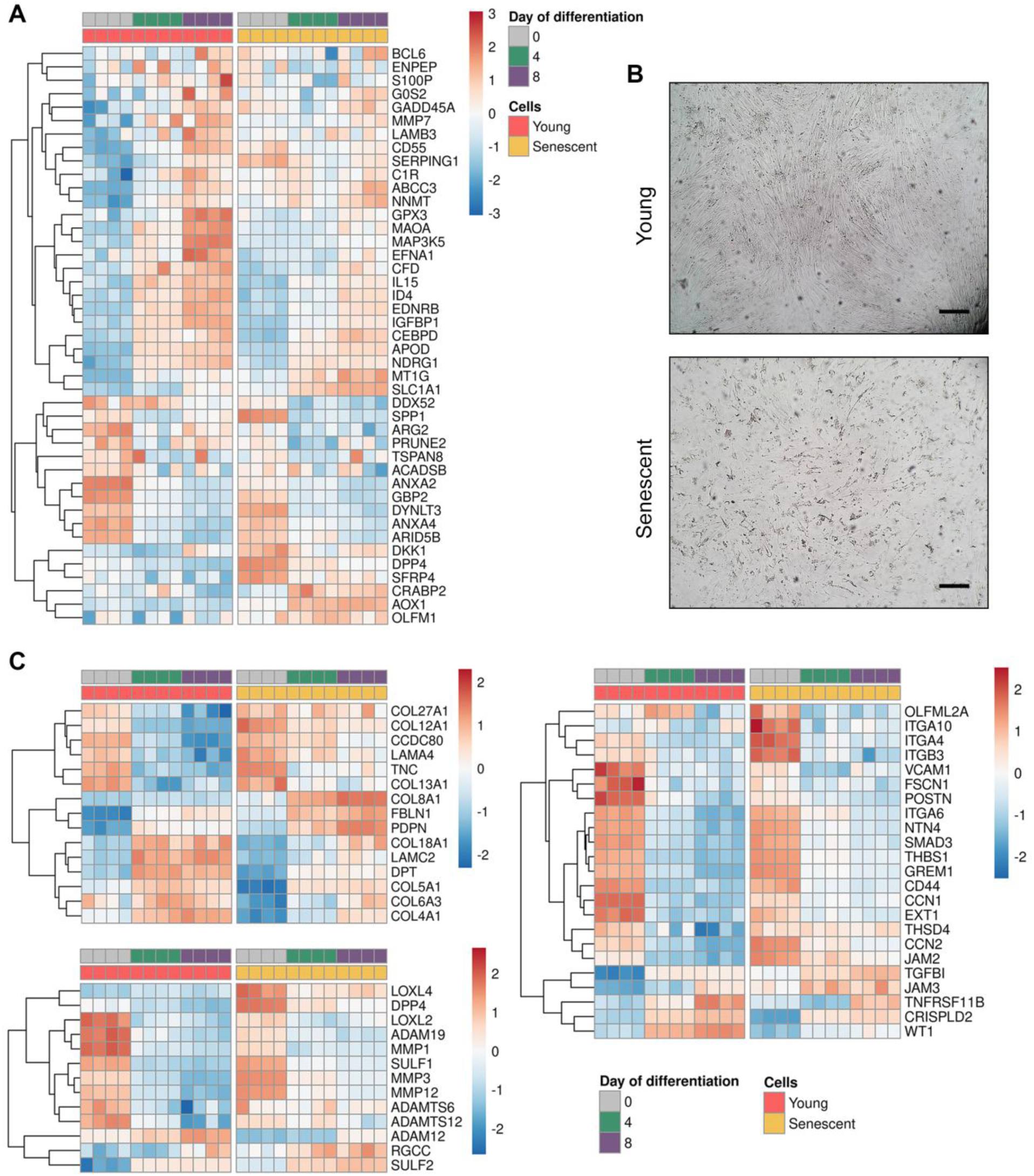
Disturbed ‘meta-signature’ of endometrial receptivity and ‘extracellular matrix organization’ upon decidualization of senescent ESCs. A Gene expression heatmap referred to “meta-signature” of endometrial receptivity for young and senescent cells during decidualization (n = 4 for each condition). B Bright-field images of ECM produced by young and senescent ESCs. Scale bars: 100 μm. C Gene expression heatmaps referred to ‘extracellular matrix organization’ GO:BP term subgrouped into ‘ECM components’, ‘ECM modifiers’ and ‘ECM regulators’ for young and senescent cells during decidualization (n = 4 for each condition). Data information: In (A) and (C), expression values are presented as VST transformed normalized counts.

Embryo implantation includes several stages: apposition (correct orientation of blastocyst towards endometrium in the uterine cavity), attachment and invasion. While the first stages mostly rely on the appropriate receptivity of endometrium, the latter stage involves direct interaction of the trophoblastic cells with decidualized stroma and its extracellular matrix (Zhu et al, 2012). According to the results of RNA-seq analysis, senescence has a huge impact on ECM organization during ESCs decidualization, what should also affect implantation process. Microphotographs presented in Fig 4B provide clear illustration of the difference in structures of ECM produced by young and senescent ESCs, namely, in case of young cells, ECM was well-organized, in case of senescent cells, it was almost completely degraded. To get better understanding of the disturbances in the ECM organization linked to ESCs senescence, we detailed ‘Extracellular matrix organization’ GO term identified above (Fig …). Normally, ESCs decidualization is accompanied by the significant reorganization of ECM. That was reflected by the distinct shifts in the expression patterns of the genes composing ECM, regulating and modifying its structure during decidual transformation of young cells (Fig 4C). Contrarily, decidualization of senescent ESCs was characterized by blurred alterations in either ECM related expression pattern, demonstrating improper ECM remodeling (Fig 4C).

Properly reorganized ECM of decidual ESCs creates so-called implantation site allowing further communication of trophoblastic cells with ESCs to form placental tissue. Cell-to-cell interaction requires expression of the specific ligands by one type of cells that would be on the ‘key-lock system’ with the receptors exposed on the other type of cells. In the study published by Pavličev with colleagues, the authors specified the pattern of ligands and receptors expressed by decidual ESCs, for which matched pairs were found in adjacent trophoblasts (Pavličev et al, 2017). When we tested the enrichment of these gene subsets among DEGs expressed by young and senescent ESCs in course of decidualization, we observed much less pronounced ‘trophoblast-interacting’ profile in case of senescent cells (FDR 3.85e-02) (Fig 5A). Together these data suggest in favor of impaired embryo implantation in presence of senescent ESCs.

**Figure 5.**
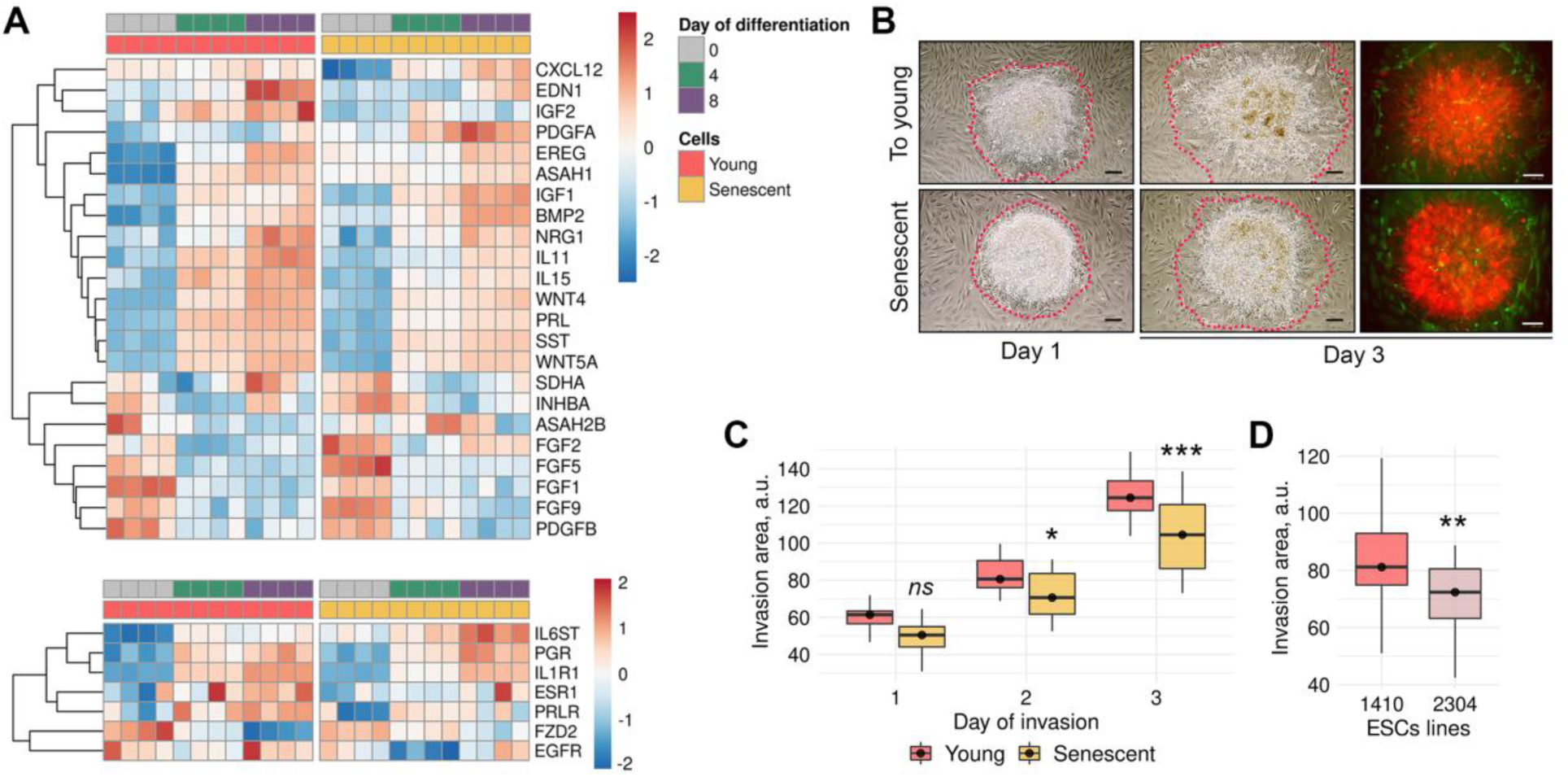
Senescent ESCs demonstrate impaired “trophoblast-interacting” profile and limited trophoblasts invasion during decidualization. A Gene expression heatmaps of core ligands (top) and receptors (bottom) responsible for interaction with trophoblasts (n = 4 for each condition). B Bright-field and immunofluorescent representative images of BeWo spheroids invasion into the monolayers of decidualizaing young or senescent ESCs. Scale bars: 100 μm. C Quantification of invasion areas of BeWo spheroids for (B) (n = 30 spheroids for each condition). D Quantification of invasion areas of BeWo spheroids into the decidualizing monolayers of ‘decidualization-prone’1410 ESCs line and of ‘senescence-prone’ 2304 ESCs line (n = 30 spheroids for each condition). Data information: In (A), expression values are presented as VST transformed normalized counts. In (C) and (D), data are presented as median ± IQR. *P<0.05, **P<0.01, ***P<0.005 (C) (two-way ANOVA with Tukey HSD), (D) (Student’s t-test).

For functional validation of the above results we used well-described ‘in vitro implantation’ model that implies estimation of the invasion area of spheroids formed from BeWo b30 cells (Grümmer et al, 1994; Weimar et al, 2013). This choriocarcinoma cell line is applied to study various aspects of implantation due to its similarity with the outer trophectoderm layer of human blastocyst (Grümmer et al, 1994). Notably, blastocyst-like spheroids demonstrated limited invasion into the monolayer of decidualizing senescent ESCs (Fig 5B and C). In order to strengthen our observations, we reproduced invasion experiments in more biologically relevant conditions, i.e. using two patients’ ESCs lines with the most pronounced differences in the severity of senescence markers and decidual response. Importantly, BeWo spheroids had much better invasion into the more ‘decidualization-prone’ line 1410, while invasion was significantly worse into the line 2304 characterized by the higher degree of senescence (Fig 5D). To sum up, we have clearly demonstrated that improper reaction of senescent ESCs towards steroid hormones negatively affects blastocyst invasion, what might lead to implantation failure.

### 5. Senescent ESCs negatively affect decidualization of their ‘healthy’ surroundings

Impaired decidualization of senescent ESCs themselves might not be the only consequence of their presence within endometrial tissue. Today it is convincingly shown that senescent cells may spread negative influence on the neighboring cells via SASP factors that can act both directly through cell contacting and distantly (Coppé et al, 2010; Borodkina et al, 2018). In our recent study we revealed that SASP factors produced by senescent ESCs caused paracrine senescence in their surroundings (Griukova et al, 2019). Here, we tested whether senescent ESCs might interfere decidualization of the neighboring young cells. To do so, we induced decidualization in the mixed cultures consisting of young ESCs expressing our genetic construct that allows estimating complex decidual response by fluorescence of the mCherry reporter and of unlabeled young or senescent ESCs. Notably, the effectiveness of decidual reaction in young ESCs was reduced in presence of senescent ones (Fig 6A). We next reproduced the same experiment in more biologically relevant 3D culture conditions. As expected, negative impact of senescent ESCs on the decidual reaction of young cells in 3D was even more pronounced than in 2D, probably, due to closer contacting (Fig 6B).

**Figure 6.**
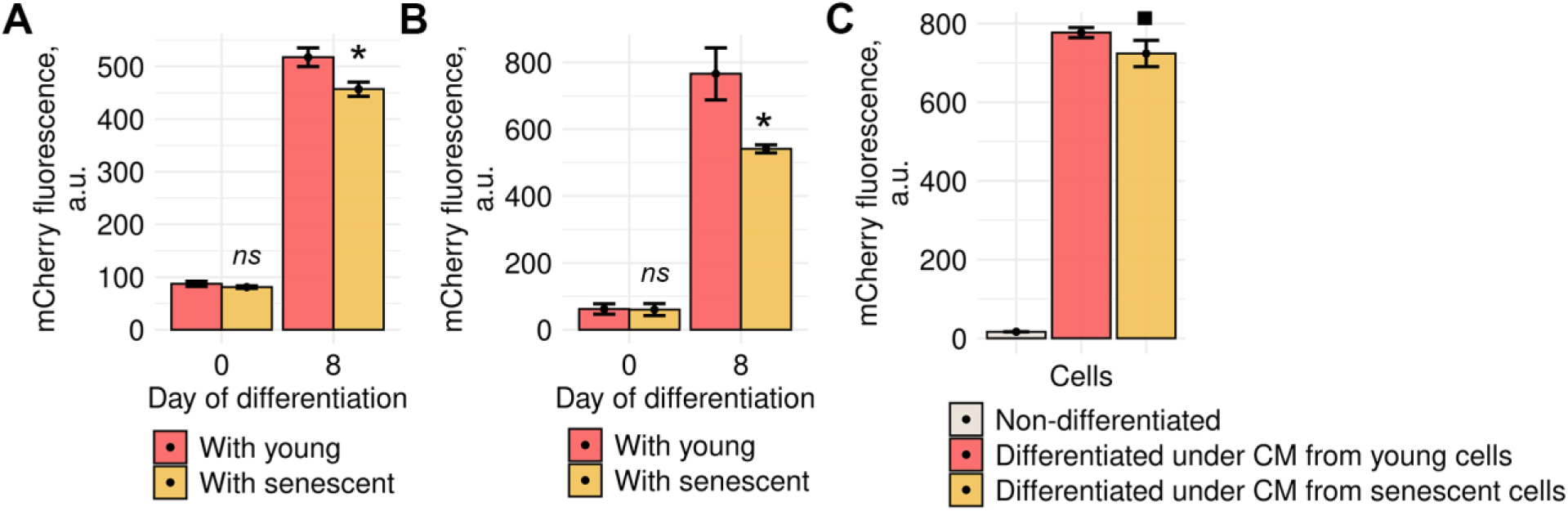
Presence of senescent ESCs causes ‘bystander’ quenching of decidual response in the nearby cells. A Fluorescence intensity of the decidual reporter protein in undifferentiated and decidualized ESCs co-cultured with young or senescent cells in 2D conditions (n = 3). B The same as for (A) in 3D conditions (n = 3). C Fluorescence intensity of the decidual reporter protein in ESCs decidualized in CM collected from young or senescent ESCs (n = 3). Data information: In (A–C), data are presented as mean ± SD. *P<0.05, ▀P<0.1 (ANOVA with Tukey HSD).

Another way of how senescent cells may influence their environment is distant action of SASP; therefore, we further tested whether conditioned media (CM) form senescent ESCs, containing soluble SASP factors, would affect decidualization of young cells. As expected, we revealed that CM from senescent ESCs diminished decidualization of control cells, though the observed effects were less pronounced than during cell co-culturing (Fig 6C).

Thereby, presence of senescent ESCs within endometrial tissue might lead to the following undesirable outcomes: (1) inability of senescent cells to respond properly to the hormonal stimulation; (2) significant reduction of the decidual reaction of the adjacent normal cells; (3) altered decidualization dynamics of normal ESCs located distantly from senescent cells. Finally, that would lead to disordered tissue remodeling during the second phase of the menstrual cycle, alterations in WOI establishment, and, probably, to impaired embryo implantation.

### 6. The impact of the crucial SASP components – reactive oxygen species (ROS) and plasminogen activator inhibitor-1 (PAI-1) – on the decidualization of young ESCs

We further tried to uncover the role of the concrete SASP components in the impairment of decidual response of the senescent cells’ surroundings. We focused on ROS and PAI-1 due to several reasons. Firstly, their role in the paracrine activity of senescent cells is well established (Coppe et al, 2010; Griukova et al, 2019; Nelson et al, 2012; Nelson et al, 2018; Vaughan et al, 2017). Secondly, both were proved to be extremely important for the proper functioning of endometrial stroma, and any alterations in time and level of their production might disturb implantation (Al-Sabbagh et al, 2011; Coulam et al, 2006; Salazar Garcia et al, 2016; Schattman et al, 2015).

ROS are small non-protein components of SASP partially responsible for the so called ‘bystander’ senescence induced in the young cells in presence of the senescent ones (Nelson et al, 2012; Nelson et al, 2018). Accumulated evidence demonstrated that adequate ROS play an integral role in initiating the endometrial decidual response, while excessive ROS can impair decidualization process and are associated with a spectrum of female reproductive disorders (Al-Sabbagh et al, 2011; Schattman et al, 2015). Previously, we revealed that senescent ESCs had increased intracellular ROS levels (Borodkina et al, 2014; Deryabin & Borodkina, in press). Here we hypothesized that due to elevated intracellular ROS levels senescent ESCs should excrete more oxidizers into the extracellular space, which may further diffuse into the neighboring young cells, shift their redox balance, thus leading to improper decidualization. Using wide-spread fluorescent probe H_2_DCF-DA, we first confirmed increased intracellular ROS levels in senescent ESCs (Fig 7A).

**Figure 7.**
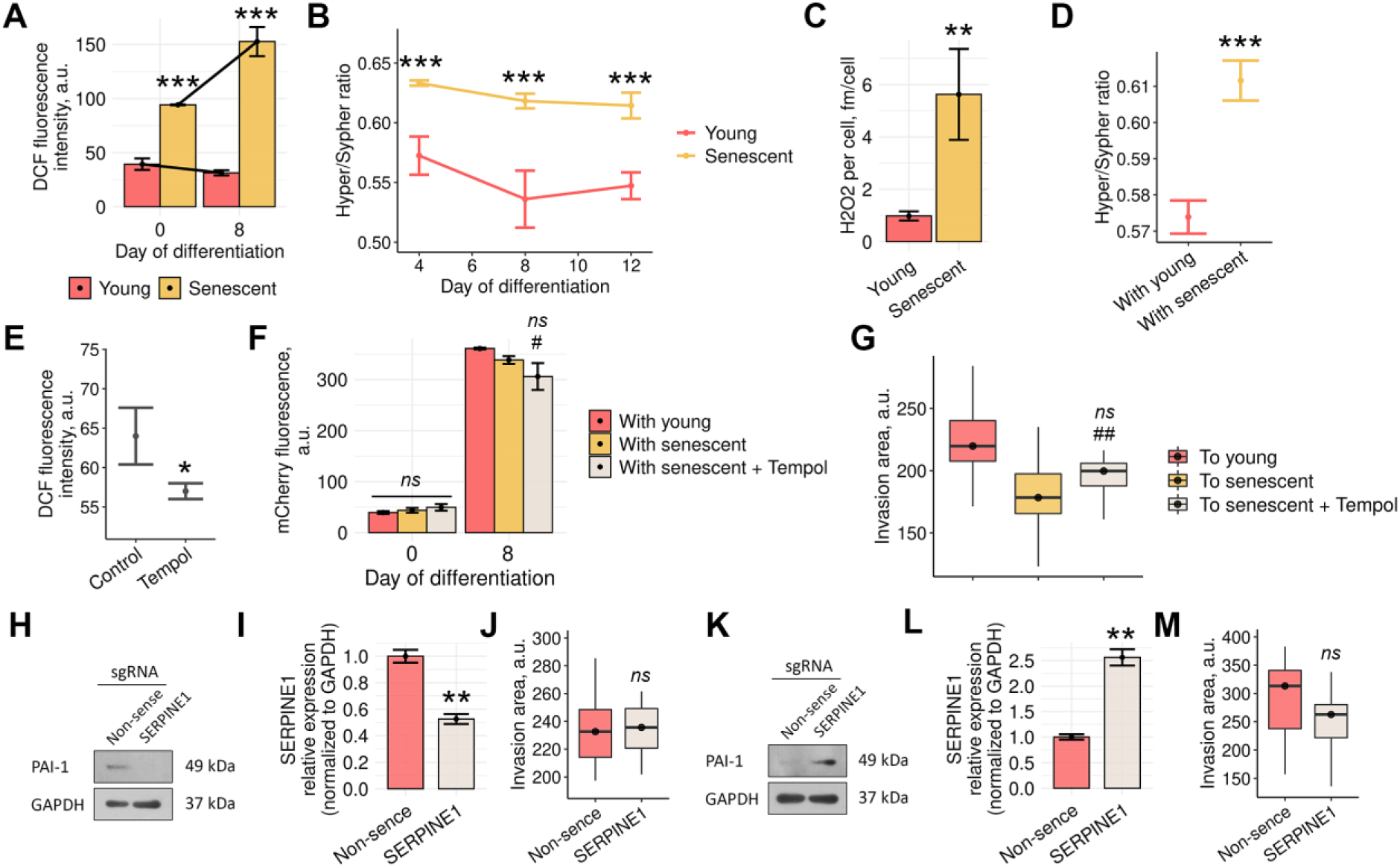
Modulating production of the crucial SASP components ROS and PAI-1 by senescent ESCs is insufficient to alleviate their negative impact on nearby cells and on trophoblasts invasion. A Intracellular ROS estimation by DCF fluorescence intensity (n = 3). B Dynamics of intracellular H_2_O_2_ levels in young and senescent ESCs in course of decidualization assessed by Hyper/SypHer ratios (n = 3). C Amounts of H_2_O_2_ excreted into the intracellular space by young and senescent ESCs (n = 3). D Hyper/SypHer ratios in Hyper/SypHer-expressing ESCs co-cultured with young or senescent cells reflecting H_2_O_2_ transmitting (n = 3). E Decreased intracellular ROS levels in ESCs upon tempol treatment estimated by DCF fluorescence intensity (n = 3). F Fluorescence intensity of the decidual reporter protein in undifferentiated and decidualized ESCs co-cultured with young, senescent, or tempol-pretreated senescent cells (n = 3). G Quantification of invasion areas of BeWo spheroids into the monolayers of decidualizaing young, senescent or tempol-pretreated senescent ESCs (n = 30 spheroids for each condition). H, I Verification of CRISPR-Cas9 mediated *SERPINE1* knockout in ESCs performed by Western blotting (H) and RT-PCR (I) (n = 3). J Quantification of invasion areas of BeWo spheroids into the monolayers of decidualizaing senescent ESCs with or without *SERPINE1* knockout (n = 30 spheroids for each condition). K, L Verification of CRISPR-Cas9 mediated *SERPINE1* overexpression in ESCs performed by Western blotting (K) and RT-PCR (L) (n = 3). M Quantification of invasion areas of BeWo spheroids into the monolayers of decidualizaing ESCs with or without *SERPINE1*overexpression (n = 30 spheroids for each condition). Data information: In (A–F), (I) and (L), data are presented as mean ± SD. In (G) and (M) data are presented as median ± IQR. *P<0.05, **P<0.01, ***P<0.005 vs senescent cells, #P<0.05, ##P<0.01 vs young cells (A), (B), (F), (D) (ANOVA with Tukey HSD), (C), (D), (E), (I), (J), (L), (M) (Student’s t-test).

In order to extend these observations, we applied genetically encoded hydrogen peroxide (H2O2) biosensor – HyPer (Belousov et al, 2006). The detailed description of this genetic construct as well as its application to estimate intracellular H_2_O_2_ levels in ESCs is provided in our previous article (Lyublinskaya et al, 2018). In brief, the intracellular level of oxidizer can be estimated by HyPer fluorescence intensity in young and senescent ESCs stably expressing HyPer. The limitation of this approach is pH-dependency of HyPer probe. This might greatly skew the results, since intracellular pH level in senescent ESCs differes significantly from that in the young cells, according to our preliminary data. To compensate for this shortcoming, we additionally applied genetically encoded pH-sensitive probe SypHer to normalize HyPer fluorescence for each tested condition. By using this approach, we were able to assess dynamic alterations in H_2_O_2_ levels during decidualization of young and senescent cells. As shown in Fig 7B, decidualization of senescent ESCs was accompanied by permanently elevated H_2_O_2_ level. Furthermore, we experimentally confirmed that senescent ESCs excreted noticeably more H_2_O_2_ into the extracellular space (Fig 7C). To answer whether H_2_O_2_ secreted by senescent ESCs might alter intracellular oxidizer’s level in the young neighboring cells, we co-cultured young HyPer-expressing cells with unlabeled senescent or young ESCs. In accordance with our hypothesis, we observed significant increase in intracellular H_2_O_2_ levels in young ESCs cultured with senescent ones, demonstrating that senescent cells can directly transmit ROS and disturb redox balance in the neighboring young cells (Fig 7D). Unexpectedly, reduction of ROS level in senescent ESCs by applying antioxidant Tempol was not able to abolish their negative impact neither on decidualization of young neighboring cells, nor on invasion of blastocyst-like spheroids (Fig 7F–G).

Another SASP component chosen was PAI-1, whose controlled expression during decidualization is required for maternal ECM remodeling and proper trophoblastic invasion (Schatz et al, 1995; Mehta et al, 2014). When analyzing SASP composition of senescent ESCs, we detected enhanced secretion of PAI-1 (Griukova et al, 2019). Within the present study we modulated expression of *SERPINE1* gene encoding for PAI-1 to reveal its possible impact on blastocyst invasion. By using CRISPR-Cas9 technology we obtained *SERPINE1* overexpressing and knockout ESCs (Fig 7H and I, Fig 7K and L). As was suggested, increased level of PAI-1 produced during decidualization of *SERPINE1* overexpressing ESCs led to decreased invasion of BeWo spheroids (Fig 7M). However, *SERPINE1* knockout did not compensate for the adverse effects of senescent ESCs on blastocyst invasion (Fig 7J).

The data obtained demonstrated that altered paracrine activity of senescent ESCs indeed may impair embryo implantation. Nevertheless, modulation of the concrete SASP components produced by senescent ESCs was not able to reverse their negative influence, suggesting for the multifactorial impact of senescent ESCs on embryo invasion.

### 7. Application of senomorphics diminished adverse effects of senescent ESCs on decidualization and implantation

Based on the above, we finally tested whether complex regulation of the secretory activity of senescent ESCs could compensate for their presence within tissue. To this end, we focused on the senomorphics – a class of drugs proved to reduce SASP secretion without affecting viability of senescent cells (Borghesan et al, 2020). We chose two compounds rapamycin and metformin, whose senomorphic activity is well established for various cell types and different models of senescence (Borghesan et al, 2020; Song et al, 2020).

We first verified that both compounds were able to reduce phenotype of senescent ESCs though did not reverse proliferation arrest, proving for their senomorphic action (Fig EV3). To test whether senomorphics were able to prevent negative influence of senescent ESCs on the decidualization of their young surroundings, we performed the set of co-culturing experiments using our in vitro models. In brief, senescent ESCs either treated or not with the chosen compounds were co-cultured with young cells stably expressing decidual reporter system. Mixed cultures of young unlabeled ESCs and young decidual reporter-expressing cells were used as the control. We induced decidualization in such mixed cultures and estimated its effectiveness by assessing fluorescence intensity of the reporter protein. It should be specifically highlighted that senescent ESCs were treated with either compound prior to decidualization. Interestingly, only metformin was able to reverse negative impact of senescent ESCs on decidual reaction of ‘healthy’ surroundings, while rapamycin had no effect (Fig 8A). We then tried to uncover the impact of senomorphics on another important consequence of ESCs senescence – impaired invasion of blastocyst-like spheroids. Treatment of senescent ESCs with metformin and rapamycin completely rescued the invasion of modeled blastocysts (Fig 8B).

**Figure 8.**
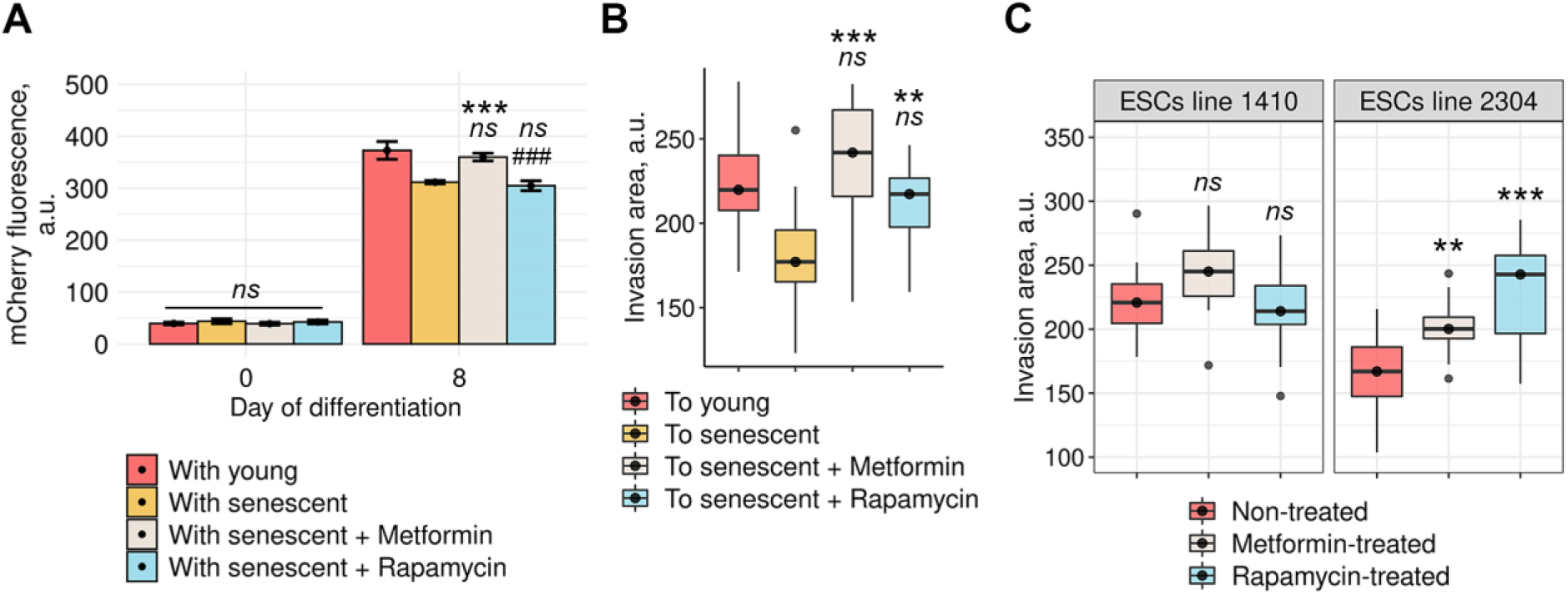
Senomorphics restore decidualization and trophoblasts invasion impaired by senescent ESCs. A Fluorescence intensity of the decidual reporter protein in undifferentiated and decidualized ESCs co-cultured with young, senescent, metformin-pretreated, or rapamycin-pretreated senescent cells (n = 3). B Quantification of invasion areas of BeWo spheroids into the monolayers of decidualizaing young, senescent, metformin-pretreated, or rapamycin-pretreated senescent ESCs (n = 30 spheroids for each condition). C Quantification of invasion areas of BeWo spheroids into the decidualizing monolayers of ‘decidualization-prone’1410 ESCs line and of ‘senescence-prone’ 2304 ESCs line either untreated or pretreated with metformin or rapamycin for 5 days prior decidualization induction (n = 30 spheroids for each condition). Data information: In (A), data are presented as mean ± SD. In (B) and (C), data are presented as median ± IQR. **P<0.01, ***P<0.005 vs senescent cells, #P<0.05 vs young cells (ANOVA with Tukey HSD).

The most intriguing part of this study was to confirm improvement of implantation upon senomorphics using patients’ ESCs lines. As described above, invasion of blastocyst-like spheroids into the monolayer of decidualizing line 1410 with minor senescence signs was significantly better compared to line 2304 characterized by the maximal degree of senescence (Fig 8C). Both ESCs lines were treated either with metformin or rapamycin for several days. After senomorphics’ removal ESCs decidualization was initiated to perform ‘in vitro implantation’ experiments. Notably, neither metformin nor rapamycin affected blastocysts invasion efficacy into ‘non-senescent’ line 1410 (Fig 8C). However, both compounds greatly enhanced invasion of blastocyst-like spheroids into the “senescent” line 2304 (Fig 8C). Such a targeted implantation improvement upon senomorphics provides additional confirmation that ESCs senescence adversely affects embryo implantation. Furthermore, application of senomorphics can be considered as effective and safe strategy to improve embryo implantation efficacy.

## Discussion

Development of cryopreservation and spreading of preimplantation genetic testing for aneuploidy (PGT-A) in IVF cycles highlighted significant role of endometrial factor for successful embryo implantation, which was underestimated previously. Transferring embryos with confirmed euploidy substantially increased positive IVF outcomes, though still not to 100%. About 30-50% of the implantation failures remained after PGT-A probably are originated from endometrial dysfunction (Altmäe et al, 2017; Tomari et al, 2020). The latter led to a completely new comprehending of the endometrium in pregnancy establishment; proper endometrial decidualization is now considered as the foundation for healthy pregnancy that creates the appropriate quality of the “soil” (Ng et al, 2020). Indeed, impaired decidualization was shown to predispose recurrent implantation failures (RIF); in particular, primary ESCs obtained from RIF patients demonstrated poor decidual reaction and limited blastocyst-like spheroid expansion (Deryabin et al, 2020; Francis et al, 2006; Salker et al, 2010). Within the present study we tested whether premature senescence of ESCs might underlie disturbed decidualization and receptivity of endometrial stroma finally mediating impaired implantation.

Before going into molecular details, we tried to reveal the link between ESCs senescence and decidualization at physiological level. To this end, we compared several primary ESCs lines obtained from patients without any endometrial pathologies planning to undergo IVF. By using this approach, we uncovered clear negative correlation between the level of senescence and the intensity of decidual reaction in the tested ESCs lines. Moreover, ESCs line with the most pronounced senescence markers demonstrated the worst implantation efficacy. These data are in line with the recent findings of Tomari with colleagues (Tomari et al, 2020). The authors analyzed expression of senescence markers (p21 and p16) in primary ESCs lines obtained from patients undergoing IVF. Based on the outcomes of IVF cycles, patients were subdivided into two groups – receptive (with successful implantation) and non-receptive (with implantation failure). Interestingly, ESCs lines expressing high levels of p21 and p16 belonged to samples from non-receptive group with failed embryo implantation. Unfortunately, no PGT-A was performed in this study to verify euploidy of the transferred embryos, what could significantly strengthen the obtained results. Nevertheless, according to our data there is an inverse correlation between the level of senescence and decidualization/implantation in primary ESCs cultures.

The most obvious way of how senescence might impair decidualization and implantation is lost proliferative capacity of senescent ESCs mediating formation of inadequate functional layer during the proliferative phase of the menstrual cycle. In accordance with this notion, previously, it was suggested that recurrent pregnancy losses (RPL) are associated with reduced plasticity of endometrial tissue due to deficiency in mesenchymal stromal cells coupled to ESCs senescence (Lucas et al, 2016). More recent study revealed the existence of highly proliferative stromal cells subpopulation – novel precursors of decidual cells – that was absent in endometrial tissue obtained from patients with RPL (Diniz-da-Costa et al, 2021). Within the present study, we tested whether improper hormonal responsiveness of senescent ESCs during the secretory phase might also contribute to the negative correlation between senescence and implantation. To test this assumption, we compared progression of decidualization in young and senescent ESCs. Indeed, we observed that senescent ESCs were unable to differentiate properly in response to hormonal stimuli. Along with the absence of clear mesenchymal-to-epithelial transition in such cells, expression and nuclear translocation of receptors of ovarian steroid hormones were also disturbed, what, in turn, mediated reduced expression and secretion of other genes and proteins crucial for decidual cells. These data verify suggestion that senescent ESCs have impaired decidualization ability. Interestingly, in earlier studies it was noted that aberrant accumulation of p21 caused by neddylation inhibition might contribute to senescence progression and impaired decidualization (Liao et al., 2015). Also, dysregulation of EPAC2 or calreticulin involved in hormonal responsiveness impaired ESCs decidualization in part through p21-mediated senescence (Kusama et al., 2014). Going further, we revealed that ESCs senescence might disturb all the properties crucial for normal functioning of endometrial stroma including communication with epithelial and endothelial cells, glands formation, vascularization and immune cells attraction. Such drastic alterations in decidual reaction of senescent ESCs led to the disordered expression of the top-priority biomarkers of receptive phase endometrium in humans, indicating for impaired receptivity. Notably, this subset of receptivity genes termed meta-signature of endometrial receptivity was proposed to be used as a diagnostic marker of receptive endometrium during infertility treatment, while alterations in this signature might be associated with pregnancy complications including implantation failure (Altmäe et al., 2017).

Since successful implantation requires bi-directional cross-talk between invading blastocyst and endometrium realized via secreted molecules, cell-to-cell or cell-to-matrix interactions, we assumed that above impairments in the decidualization of senescent ESCs detected by RNA-seq should necessarily affect implantation. Indeed, upon decidualization senescent ESCs demonstrated disorganized ECM architecture, due to deregulated expression of ECM components, regulators and modifiers, and disturbed expression of ligands and receptors responsible for interaction with trophoblasts. Together that hindered blastocysts’ invasion as was verified by the limited invasion of blastocyst-like spheroids into the monolayers of decidualizing senescent ESCs and of decidualizing 2304 ESCs line characterized by the pronounced senescence phenotype. In line with these results, earlier it was shown that primary ESCs obtained from RIF patients characterized by reduced decidualization also demonstrated impaired implantation of modeled spheroids (Huang et al, 2017).

Along with the proliferation block and diminished differentiation capacity, altered secretory profile of senescent cells is regarded to be responsible for the undesirable outcomes of their presence within tissues. SASP produced by senescent cells creates chronic pro-inflammatory microenvironment, alters ECM organization and may mediate senescence spreading via paracrine action on the neighboring cells (Coppé et al, 2010; Griukova et al, 2019). In our recent study we analyzed concrete composition of SASP secreted by senescent ESCs and proved for the existence SASP-mediated paracrine senescence in the young cells (Griukova et al, 2020). Here we tried to uncover whether the paracrine activity of senescent ESCs might disturb decidual response of the young neighboring cells. Indeed, we observed that both the presence of senescent ESCs and conditioned media containing SASP factors significantly reduced the efficacy of decidualization in the young ones, probably, via the secreted factors that can act both distantly and directly through cell contacts. Thus, paracrine activity of senescent ESCs should exaggerate their negative role in the functioning of endometrial tissue. Several literary findings favor this suggestion. It was shown that primary cultures obtained from RPL patients displayed prolonged proinflammatory secretory profile upon decidualization (Macklon & Brosens, 2014; Salker et al, 2012). In line with these data, Lucas et al. speculated that premature senescence and tissue inflammation might predispose RPL (Lucas et al, 2016a). Another study performed unbiased secretome analyzes of primary ESCs lines obtained from patients undergoing IVF (Durairaj et al, 2017). According to further IVF outcomes, the authors revealed that in case of successful implantation pro-inflammatory secretory profile of ESCs was strictly regulated in time, while in case of failed implantation it was prolonged and highly disordered, with the absence of clear peaks, in many ways similar to SASP. Based on that, authors suggested that deficient or damaged progenitor ESCs might be associated with RIF.

Among various factors secreted by senescent cells, ROS are of extreme importance in the context of the present study. Accumulated evidence demonstrate that ROS play integral role in ESCs decidual response, while excessive ROS and oxidative stress are liked to impaired decidualization and various female reproductive disorders including RPL (Al-Sabbagh et al, 2011; Schattman et al, 2015). Being important non-protein SASP components, ROS produced by senescent cells were recently shown to shift redox balance in the young neighboring cells (Nelson et al, 2012; Nelson et al, 2018). According to our data, senescent ESCs had elevated intracellular ROS levels throughout decidualization and displayed enhanced secretion of H2O2 into the extracellular space. Moreover, using rather elegant approach based on genetically encoded hydrogen peroxide sensor, we confirmed that senescent ESCs can directly transmit oxidizers to the young neighboring cells. Despite these results, application of antioxidant Tempol was not able to reverse negative influence of senescent ESCs on the decidualization of young surroundings as well as on the efficacy of attachment and invasion of the modeled blastocysts.

Another well-known SASP component that might have impact on the implantation process is PAI-1. Previously, we revealed that senescent ESCs secrete increased amounts of PAI-1, which is responsible for senescence transmitting (Griukova et al, 2019). With regard to implantation, waves of PAI-1 secretion should be strictly regulated in time, as an accurate balance between coagulation and fibrinolysis is mandatory for trophoblastic invasion (Mehta et al, 2014). Any disturbances in PAI-1 level can lead to pregnancy complications (Salazar Garcia et al, 2016). Particularly, increased level of PAI-1 limits trophoblastic invasion and is considered to be a risk-factor for implantation failure (Coulam et al, 2006). In line with this notion, we observed that invasion of blastocyst-like spheroids into the monolayer of decidualizing ESCs overexpressing PAI-1 was impaired. Nevertheless, PAI-1 knockout did not improve invasion of the modeled blastocysts.

The data described above evidenced that SASP is an additional mechanism of how senescent ESCs might disturb decidualization and implantation. However, removal of the individual SASP components (e.g. ROS or PAI-1) is insufficient to neutralize negative effects of senescent ESCs, what evidences for the multifactorial action of SASP. Thus, we finally tested whether complete SASP prevention by the senomorphics, namely rapamycin and metformin, could abolish adverse influence of senescent cells on decidualization of their “healthy” surroundings as well as on blastocyst implantation. Interestingly, among the compounds applied only metformin was able to prevent both undesirable outcomes of ESCs senescence. Treatment of senescent cells with rapamycin positively affected invasion of spheroids. Such differences might be due to varied effectiveness of the chosen senomorphics to suppress SASP production by senescent ESCs. Altogether, these findings contradict to some extent the existing literary data. Namely, it was shown that application of rapamycin led to decreased expression of PRL and IGFBP1, indicating impaired decidualization (Brighton et al, 2017). Another senomorphic compound – resveratrol – was also shown to have anti-deciduogenic properties reducing expression of PRL and IGFBP1 (Ochiai et al, 2019a). Moreover, the results of the clinical study demonstrated that continuous supplementation of resveratrol led to lower implantation rates (Ochiai et al, 2019b). Contrarily, we revealed that application of rapamycin or metformin improved ability of ESCs line with impaired decidual reaction (probably due to enhanced level of senescence) to accept blastocysts. The recently published study provided probable experimental explanation for this disagreement. The authors demonstrated that negative or beneficial outcomes of resveratrol application are strictly related on the phase of the menstrual cycle (Kuroda et al, 2020). If resveratrol supplementation is restricted to proliferative phase, similar to what we performed in the present study using other senomorphics, it would not adversely impact on embryo implantation or ESCs decidualization (Kuroda et al, 2020). However, when this compound is added during WOI it inhibits decidual transformation of the endometrium, what was discovered in the mentioned above studies (Brighton et al, 2017; Ochiai et al, 2019a).

Taken together, presence of senescent ESCs within endometrium as well as SASP secretion reduce hormonal responsiveness of this tissue and impair embryo implantation. Application of senomorphics during proliferative phase of the menstrual cycle may be considered as the promising strategy to reduce SASP production and to prepare tissue for further embryo implantation.

## Materials and Methods

### 1. Endometrial sampling and primary ESCs cultures

The study was approved by the Local Bioethics Committee of the Institute of Cytology of the Russian Academy of Sciences. Endometrial biopsies were obtained under the cooperation agreement with the Almazov National Medical Research Centre. None of the subjects were on hormonal treatments prior to the procedure. Written informed consents were obtained from all participants in accordance with the guidelines in The Declaration of Helsinki 2000. ESCs were isolated from endometrial tissues as described previously (Deryabin et al, 2021). Cells were cultured in DMEM/F12 (Gibco BRL, USA) at 37 °C in humidified incubator, containing 5 % CO_2_. Cultural media was supplemented with 10 % FBS (HyClone, USA), 1 % penicillin-streptomycin (Gibco BRL, USA) and 1 % glutamax (Gibco BRL, USA). Serial passaging was performed when the cells reached 80%–90% confluence.

### 2. BeWo b30 culture conditions

The original BeWo cell line is available through American Type Culture Collection (ATCC; cat. no. CCL-98). More adhesive BeWo clone (b30) used in the present study was created by Dr. Alan Schwartz (Washington University, St. Louis, MO). BeWo b30 were obtained from Scientific Research Centre Bioclinicum (Moscow, Russia) by agreement with Dr. Alan Schwartz (Washington University, St. Louis, MO). Cells were cultured in DMEM/F12 (Gibco BRL, USA) at 37 °C in humidified incubator, containing 5 % CO_2_. Cultural media was supplemented with 10 % FBS (HyClone, USA), 1 % penicillin-streptomycin (Gibco BRL, USA) and 1 % glutamax (Gibco BRL, USA).

### 3. Decidualization induction

Confluent ESCs monolayers were decidualized in DMEM/F12 containing 2 % FBS supplemented with 0.3 mM N6,2′-O-Dibutyryladenosine 3′,5′-cyclic monophosphate sodium salt (cAMP) (Sigma-Aldrich, USA), 10 nM β-Estradiol (E2) (Sigma-Aldrich, USA) and 1 µM medroxyprogesterone 17-acetate (MPA) (Sigma-Aldrich, USA).

### 4. Senescence induction and conditioned media collection

For oxidative stress-induced senescence ESCs were treated with 200 µM H_2_O_2_ (Sigma-Aldrich, USA) for 1 h. Cells were considered senescent not earlier than 14 days after treatment. ESCs were considered replicatively senescent after 25 passages. CM was collected according to the procedure described in detail in Griukova et al, 2019.

### 5. Flow cytometry

Measurements of proliferation, cell size, autofluorescence (lipofuscin accumulation), the intensity of decidual reaction, levels of intracellular ROS and H_2_O_2_ were carried out by flow cytometry. Flow cytometry was performed using the CytoFLEX (Beckman Coulter, USA) and the obtained data were analyzed using CytExpert software version 2.0. Adherent cells were rinsed twice with PBS and harvested by trypsinization. Detached cells were pooled and resuspended in fresh medium and then counted and analyzed for autofluorescence. The cell size was evaluated by cytometric forward light scattering. Intracellular ROS levels were assessed using H_2_DCF-DA dye (Invitrogen, USA). The staining procedure was carried out as described previously (Griukova et al, 2019). For estimation of decidual response ESCs transduced by the Dec_pPRL-mCherry lentiviruses were used. The detailed procedure is described in our recent study (Deryabin et al., 2021). To assess intracellular H_2_O_2_/pH levels HyPer/SypHer-expressing cells were used. The technical features of these cytometric measurements can be found in the previous studies (Lyublinskaya et al., 2018; Deryabin & Borodkina, in press).

### 6. Western blotting

Western blotting was performed as described previously (Borodkina et al., 2014). SDS-PAGE electrophoresis, transfer to nitrocellulose membrane and immunoblotting with ECL (Thermo Scientific, USA) detection were performed according to standard manufacturer’s protocols (Bio-Rad Laboratories, USA). Antibodies against the following proteins were used: glyceraldehyde-3-phosphate dehydrogenase (GAPDH) (clone 14C10) (#2118, Cell Signaling, USA), E-cadherin (clone HECD-1) (ab1416, Abcam, UK), vimentin (clone RV202) (ab8978, Abcam, UK), progesterone receptor A/B (clone D8Q2J) (#8757, Cell Signaling, USA), estrogen receptor α (clone D6R2W), HMGB1 (clone D3E5) (#6893, Cell Signaling, USA), phospho-p53 (Ser15) (clone 16G8) (#9286, Cell Signaling, USA), p21 (clone 12D1) (#2947, Cell Signaling, USA), phospho-Rb (Ser807/811) (#8516, Cell Signaling, USA), p16 INK4A (clone D3W8G) (#92803, Cell Signaling, USA), PAI-1 (clone D9C4) (#11907, Cell Signaling, USA) as well as horseradish peroxidase-conjugated goat anti-rabbit IgG (GAR-HRP, Cell Signaling, USA) and anti-mouse IgG (GAM-HRP, Cell Signaling, USA).

### 7. Senescence-associated β-galactosidase staining

Senescence-associated β-galactosidase staining was performed using senescence β-galactosidase staining kit (Cell Signaling, USA) according to manufacturer’s instructions. Quantitative analysis of images was produced with the application of MatLab package, according to the algorithm described in the relevant paper (Shlush et al, 2011). For each experimental point not less 50 randomly selected cells were analyzed.

### 8. RNA extraction, reverse transcription and real time PCR

RNA extraction, reverse transcription and real time PCR were performed as described in our previous study (Griukova et al, 2019). Reagents for RNA extraction (ExtractRNA reagent), for reverse transcription (MMLV RT kit) and for real time PCR (HS SYBR kit) were obtained from Evrogen, Russia. Gene expression levels were assessed using the Realtime PCR BioRad CFX-96 amplifier (BioRad, USA), the following analysis of the obtained data was performed using the Bio-Rad CFX Manager software (BioRad, USA). Primer sequences and the corresponding annealing temperatures are listed in Table 2.

**Table 2.**
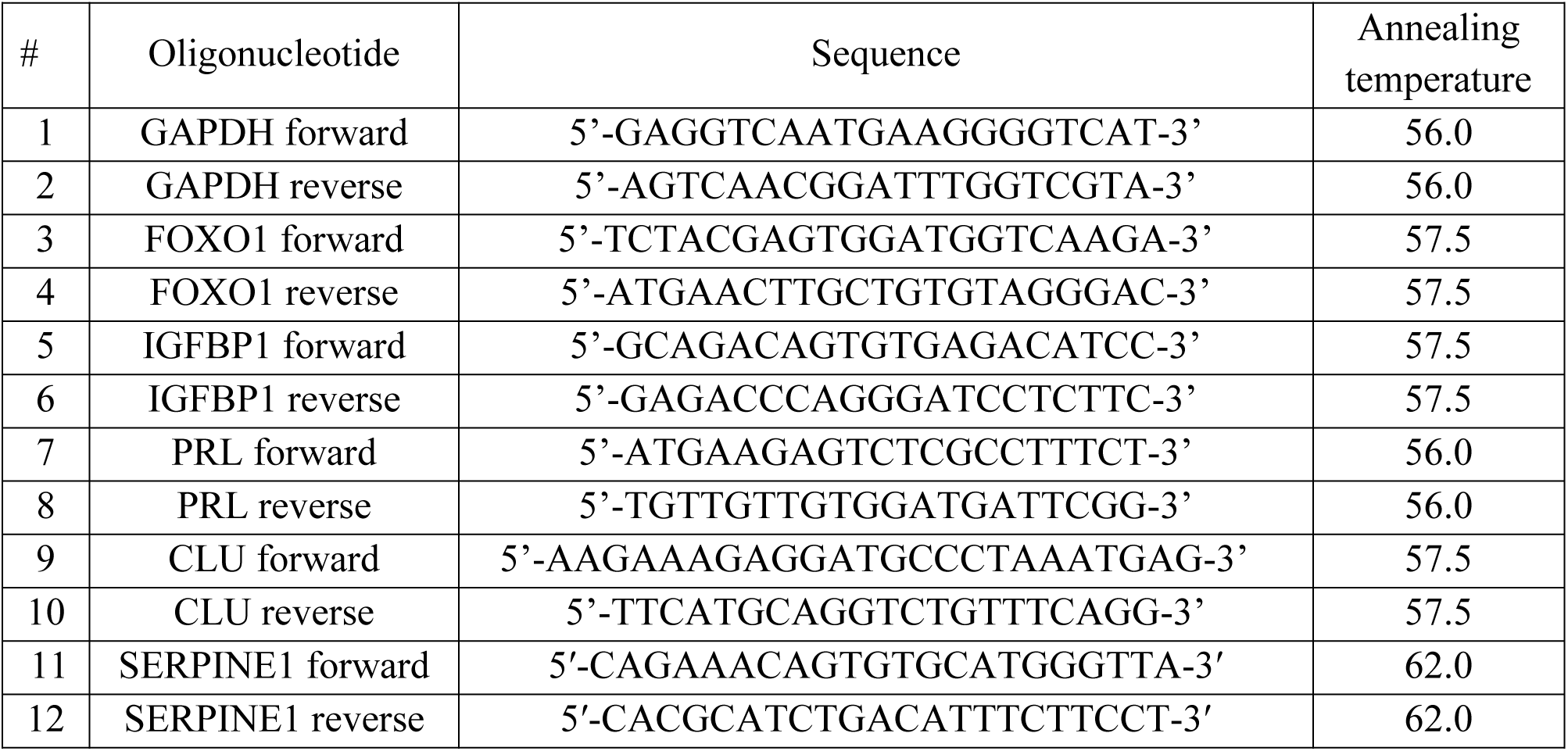
Primer oligonucleotide sequences.

### 9. Molecular cloning

For CRISPR-mediated *SERPINE1* knockout and transactivation the following lentivectors were used: pCC_01 - hU6-BsmBI-sgRNA(E+F)-barcode-EFS-Cas9-NLS-2A-Puro-WPRE (Addgene 139086, Legut et al, 2020) and pCC_05 - hU6-BsmBI-sgRNA(E+F)-barcode-EFS-dCas9-NLS-VPR-2A-Puro-WPRE (Addgene 139090, Legut et al, 2020). Oligonucleotide sequences for single guide RNAs (sgRNAs), sgRNAs design and cloning procedures were performed in accordance with previously described (Deryabin et al, 2019).

### 10. Lentiviral transduction

ESCs were transduced with the following lentiviruses: Dec_pPRL-mCherry (Deryabin et al., 2021), cyto-HyPer and SyPher (Lyublinskaya et al., 2018; Deryabin & Borodkina, in press), CRISPR-Cas9-based lentivuruses for SERPINE1 knockout or overexpression. Protocols of lentiviral particles production and ESCs lentiviral transduction are described in detail in our previous article (Deryabin et al., 2019).

### 11. ELISA

The amounts of secreted prolactin were quantified in the cell supernatants by the Prolactin Human ELISA Kit (Abcam, USA). The data were normalized to the total amount of protein determined by the Bradford method. To determine the concentration of secreted proteins in samples, GraphPad Prism 5 was used.

### 12. Immunofluorescence

Cells grown on coverslips were fixed with 4 % formaldehyde (15 min), permeabilized with 0.1 % Triton X-100 (10 min) and blocked with 1% BSA (1 h). Cells were incubated with progesterone receptor A/B (clone D8Q2J) (#8757, Cell Signaling, USA) primary antibodies overnight at 4 °C, followed by the incubation with secondary antibodies ‒ Alexa Fluor 568 goat anti-rabbit (Invitrogen, USA) for 1 h at room temperature. The slides were counterstained with 1 μg/ml DAPI (Sigma-Aldrich, USA), mounted using 2 % propyl gallate and analyzed using Olympus FV3000 confocal microscope (Olympus, Japan). To visualize F-actin cytoskeleton, cells were fixed, permiabilized and blocked as described above and then incubated with rhodamine phalloidin (Thermo Scientific, USA) for 30 min at 37°C. ZOE Fluorescent Cell Imager (BioRad, USA) was used to view and acquire images.

### 13. Bioinformatics

RNA-seq reads processing, lightweight-mapping, transcript abundances estimation, and differential expression analysis were conducted as described previously (Deryabin et al, 2021). Summarized to a gene level expression count matrix was filtered to contain rows having at least 2 estimated counts across half of the samples, the resulting matrix contained 19609 genes. Differential expression analysis and log fold changes (LFC) estimation were computed using DESeq2 (version 1.26.0) with the use of LRT-test (Love et al, 2014) (Table 1) with reducing formula to get difference for interaction between senescence state and differentiation status. 2963 genes with estimated FDR < 0.01 were defined to be statistically differentially expressed and subjected further to co-expession analysis. Clusterization of identified DEGs were performed with the use of DEGreport package (version 1.28.0). Obtained clusters were further represented via heatmaps and functionally annotated in GO:BP terms using pheatmap (version 1.0.12) and clusterProfiler (version 3.14.3) R packages as described previously (Yu et al, 2012; Deryabin et al, 2021).

### 14. Preparation of spheroids from ESCs or BeWo b30

Spheroids were formed using the hanging drop technique. 7*10^3^ cells per 35 μL were placed in drops on the cover of 100 mm culture dishes and then inverted over the dish. For effective formation of BeWo b30 spheroids 0.2 % methylcellulose was added. Cells spontaneously aggregated in hanging drops for 48 h and then were transferred into dishes coated with 2-hydroxyethyl methacrylate (HEMA; Sigma-Aldrich, USA). Single cell suspension was obtained by ESCs spheroid treatment with 0.05 % trypsin/EDTA to assess the effectiveness of ESCs decidualization.

### 15. In vitro invasion model

Spheroids formed from BeWo b30 were seeded on top of the monolayers of decidualized ESCs. Cells were cultured for 72 h (except for Fig 5B and C) at 37 °C in humidified incubator, containing 5 % CO_2_ before imaging. Quantitative analysis of the invasion area was produced with the application of ImageJ software. Not less than 30 spheroids for each sample were analyzed.

### 16. Estimation of H_2_O_2_ excretion

To assess the amounts of H_2_O_2_ excreted by ESCs into the extracellular space Amplex™ Red Hydrogen Peroxide/Peroxidase Assay Kit (Invitrogen, USA) was used according to manufacturer’s instructions.

### 17. Antioxidant and senomorphics supplementation schemes

Senescent ESCs were treated with 2 mM tempol (Santa Cruz Biotechnology, USA), 200 nM rapamycin (Calbiochem, USA), 5 mM metformin (Merk, Germany) for 7 days prior to decidualization induction. Cell culture medium supplemented with either compound was changed daily.

### 18. Statistical analysis

All quantitative data are shown as mean ± SD or as median ± IQR (indicated for each figure). To get significance in the difference between two groups Students t-test was applied. For multiple comparisons between groups, one- or two-way ANOVA with Tukey HSD was used. Statistical analysis was performed using R software version 4.1.

## Data availability

RNA-Seq data: Gene Expression Omnibus GSE160702 (https://www-ncbi-nlm-nih-gov.ezproxy.u-pec.fr/geo/query/acc.cgi?acc=GSE160702)

The raw data generated during and/or analyzed in the current study are available from the corresponding author on request.

## Acknowledgements

This study was funded by the Russian Science Foundation (grant number: 19-74-10038). The authors are thankful to Dr. Alla Shatrova for the assistance in the flow cytometry analysis, to Dr. Olga Lyublinskaya for the assistance in HyPer/SyPher assessment, and to Maria Sirotkina for the assistance in the figures design.

## Author contributions

AVB supervised the work, wrote and edited the manuscript. AVB and PID designed the study and performed most of the experiments. PID in designed and conducted bioinformatic analysis, performed statistical analysis of the obtained data. YSI performed HyPer/SypHer measurements.

## The Paper Explained

### PROBLEM

Infertility is a crucial health issue worldwide. The number of women of reproductive age suffering from infertility increases annually. In vitro fertilization (IVF) is considered to be the leading approach for infertility curing. Transferring embryos with confirmed euploidy due to preimplantation genetic testing (PGT-A) for aneuploidy substantially increased IVF efficacy, though still not to 100 %. Therefore, the search for new approaches to improve the successful outcomes of IVF is ongoing. About one-third of implantation failures remained after PGT-A are regarded to be mediated by inadequate endometrial receptivity. Particularly, impaired decidualization (hormone-regulated differentiation of endometrial stromal cells into decidual ones) is proved to mediate implantation failures. However, molecular mechanisms underlying disturbed decidualization as well as the ways to improve it are not yet clear.

## RESULTS

We found that premature senescence of endometrial stromal cells (ESCs) might be the cause for impaired decidualization. Using both in vitro models and patients’ lines we revealed that senescence negatively affected hormone-induced decidual transformation of the stromal compartment of endometrium. Application of bioinformatics uncovered crucial disturbances in decidual reaction of senescent ESCs that might affect embryo invasion, such as altered “meta-signature” of human endometrial receptivity, disturbed ligand-receptor interaction with trophoblasts and modified architecture of extracellular matrix. These bioinformatic predictions of impaired embryo implantation were functionally validated using in vitro implantation model. Moreover, we observed that senescent ESCs probably via altered secretome caused “bystander” quenching of decidual reaction in adjacent cells reinforcing dysfunction of stromal compartment. Implementation of senomorphics that reduced senescence phenotype diminished adverse effects of senescent ESCs on decidualization and implantation in both in vitro models and patients’ lines.

## IMPACT

Presence of senescent ESCs within stromal compartment of the endometrium might be considered as a risk-factor for embryo implantation failure. Application of senomorphics during the proliferative phase of the menstrual cycle seems to be a promising strategy to alleviate negative effects of senescent ESCs and to increase implantation rates during IVF treatment.

---

**Expanded View Figure 1.**
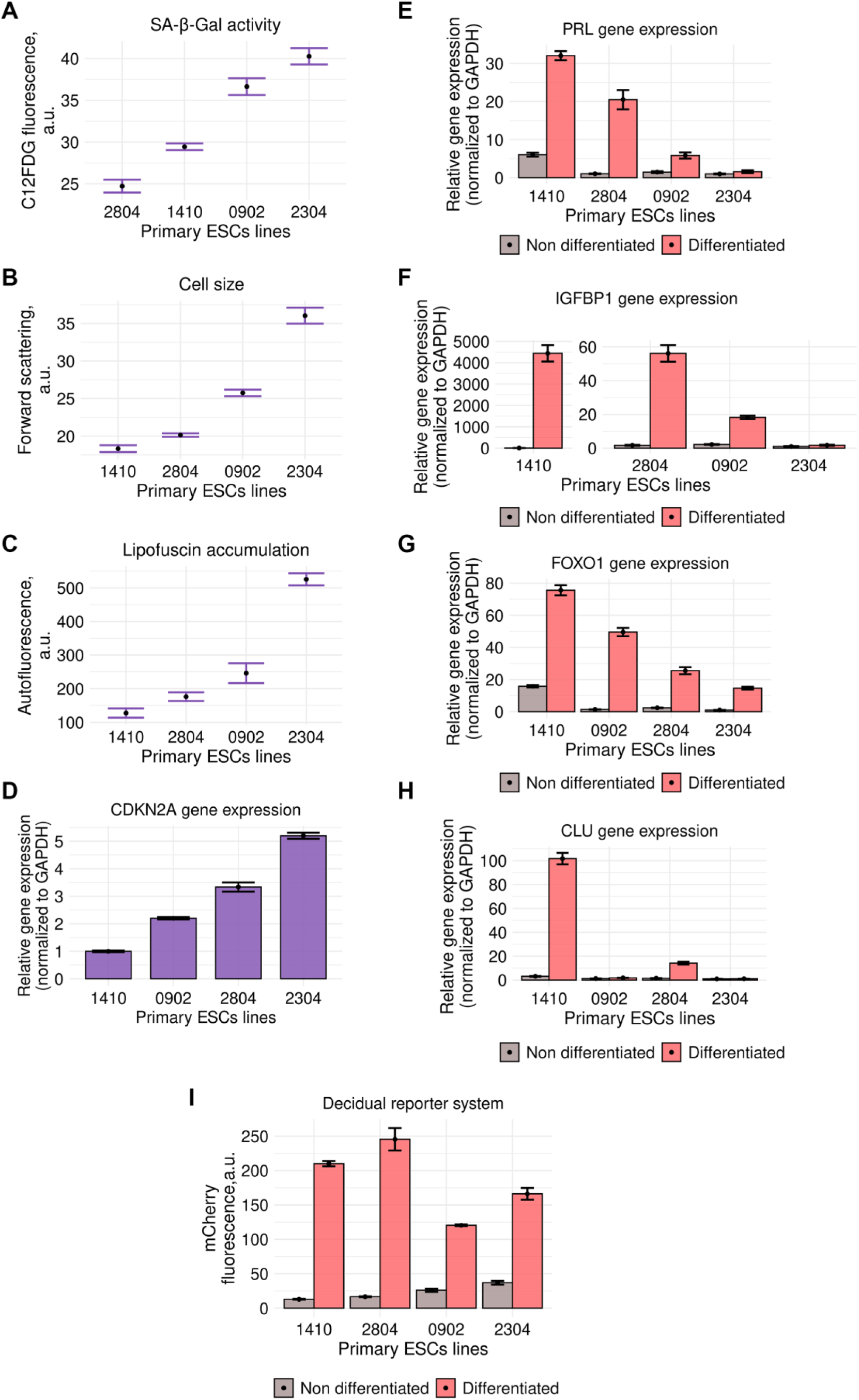
Senescence and decidual markers assessed in primary ESCs cultures. A, B, C, D Assessment of senescence markers in primary ESCs lines: SA-β-Gal activity (A), cell size (B), lipofuscin accumulation (C), expression of *CDKN2A*gene (D) (n = 3 for each sample). E, F, G, H, Estimation of the core parameters of decidual differentiation in primary ESCs lines: expression of *FOXO1* (E), *PRL* (F), *IGFBP-1* (G), *CLU* (H) genes and fluorescence intensity of decidual reporter protein (I) (n = 3 for each sample). Data information: Data are presented as mean ± SD.

**Expanded View Figure 2.**
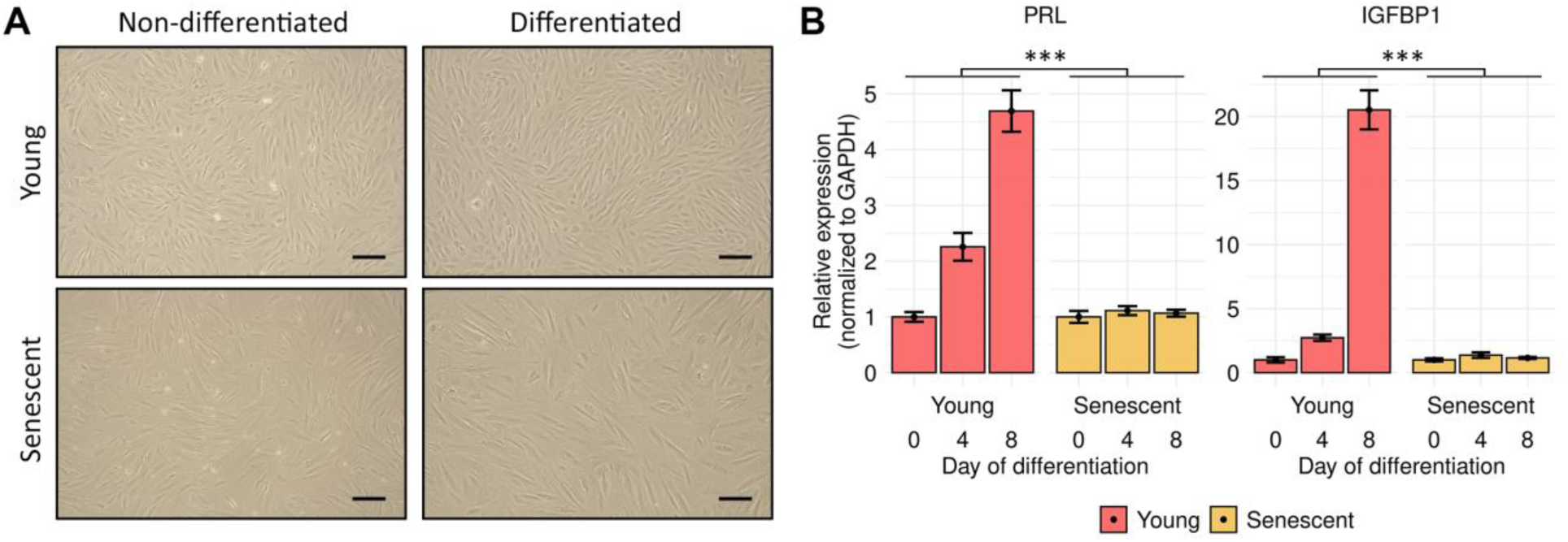
Impaired decidualization of replicatively senescent ESCs. A Bright-field images of young and replicatively senescent ESCs before and after decidualization. Scale bars: 100 μm. B Quantitative RT-PCR detection of *PRL* and *IGFBP1* expression levels in young and replicatively senescent ESCs (n = 3 for each sample). Data information: Data are presented as mean ± SD. ***P<0.005 (two-sided ANOVA).

**Expanded View Figure 3.**
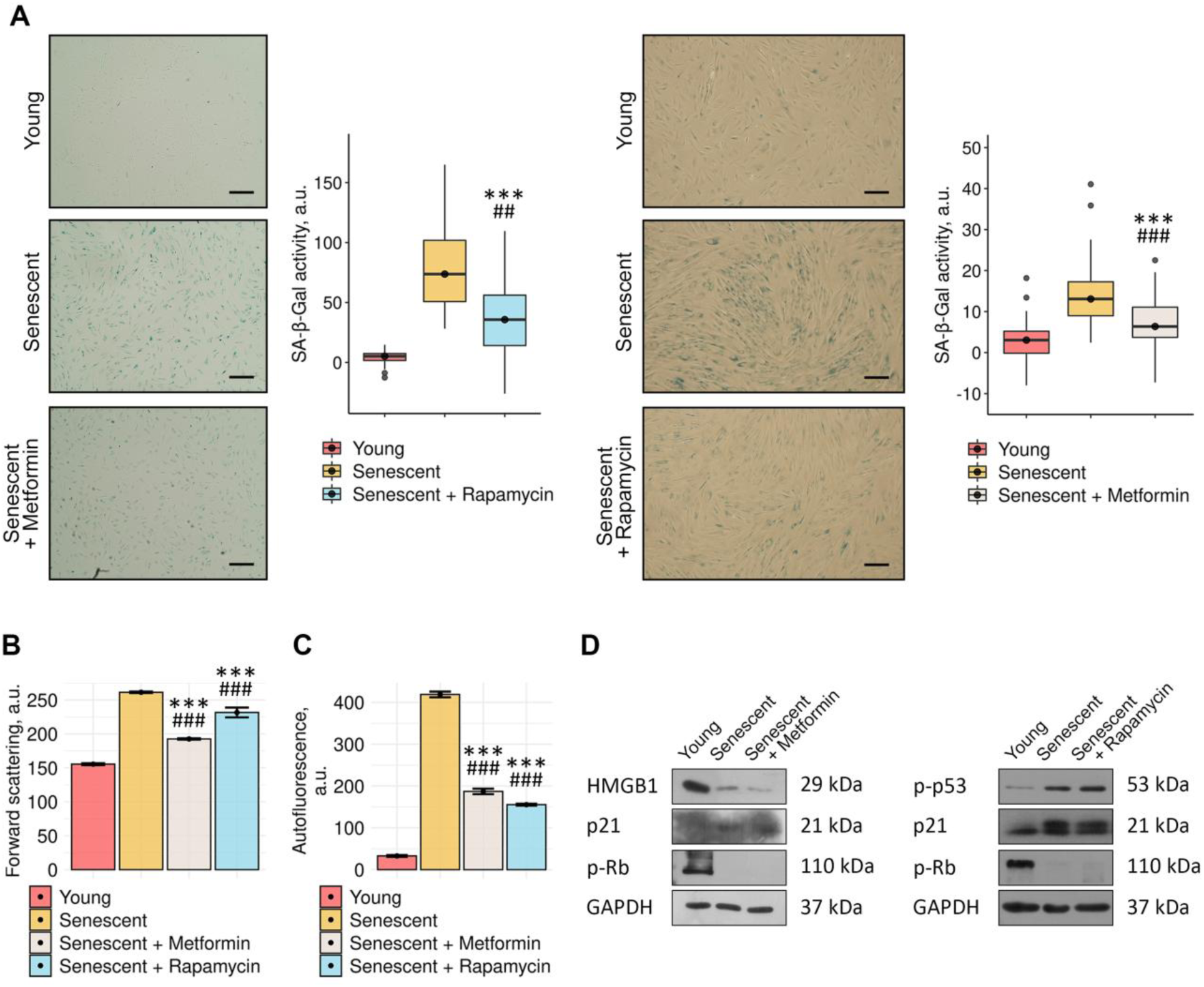
Senomorphic compounds – rapamycin and metformin – reduce phenotype of senescent ESCs. A Representative images and quantification of SA-β-Gal activity in young, senescent and rapamycin/metformin-pretreated senescent ESCs (n = 50). B, C, D Assessment of cell size (B), lipofuscin accumulation (C), expression levels of HMGB1, p21 and phosphorylation levels of p53 and Rb proteins (D) in young, senescent and rapamycin/metformin-pretreated senescent ESCs (n = 3). Data information: In (A), data are presented as median ± IQR. In (B) and (C), data are presented as mean ± SD. ***P<0.005 vs senescent cells, ##P<0.01, ###P<0.005 vs young cells (ANOVA with Tukey HSD).

## Appendix

**Appendix FigS1–22 – The multifaceted analysis of each cluster presented in** **Fig 3B** **as well as of the non-clustered DEGs.**

---

**Appendix Fig S1.**
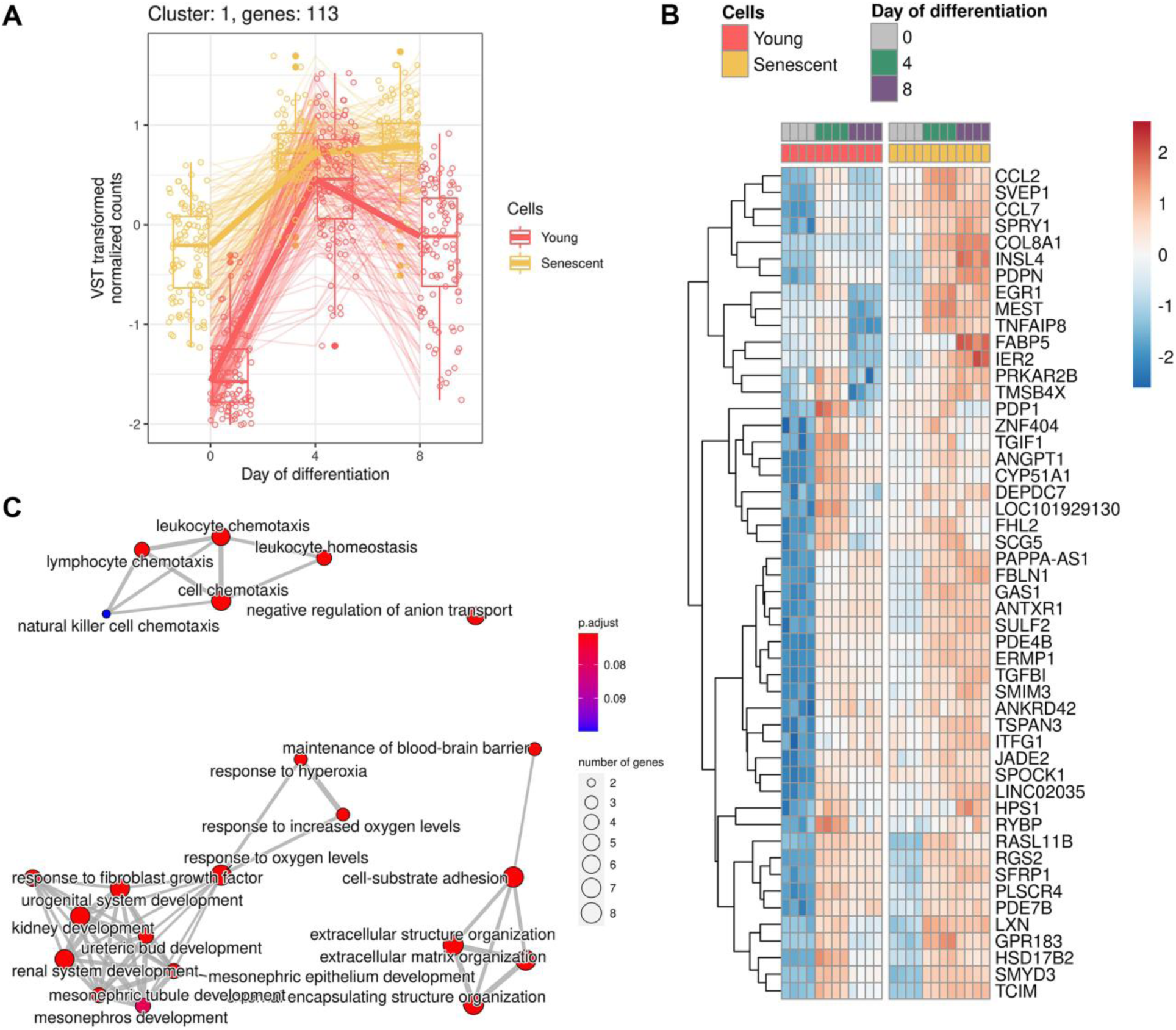
Cluster 1. A Expression of genes forming cluster B Heatmap reflecting expression of the top 50 DEGs in cluster C Functional enrichment analysis (FEA) of clustered genes in GO:BP terms

**Appendix Fig S2.**
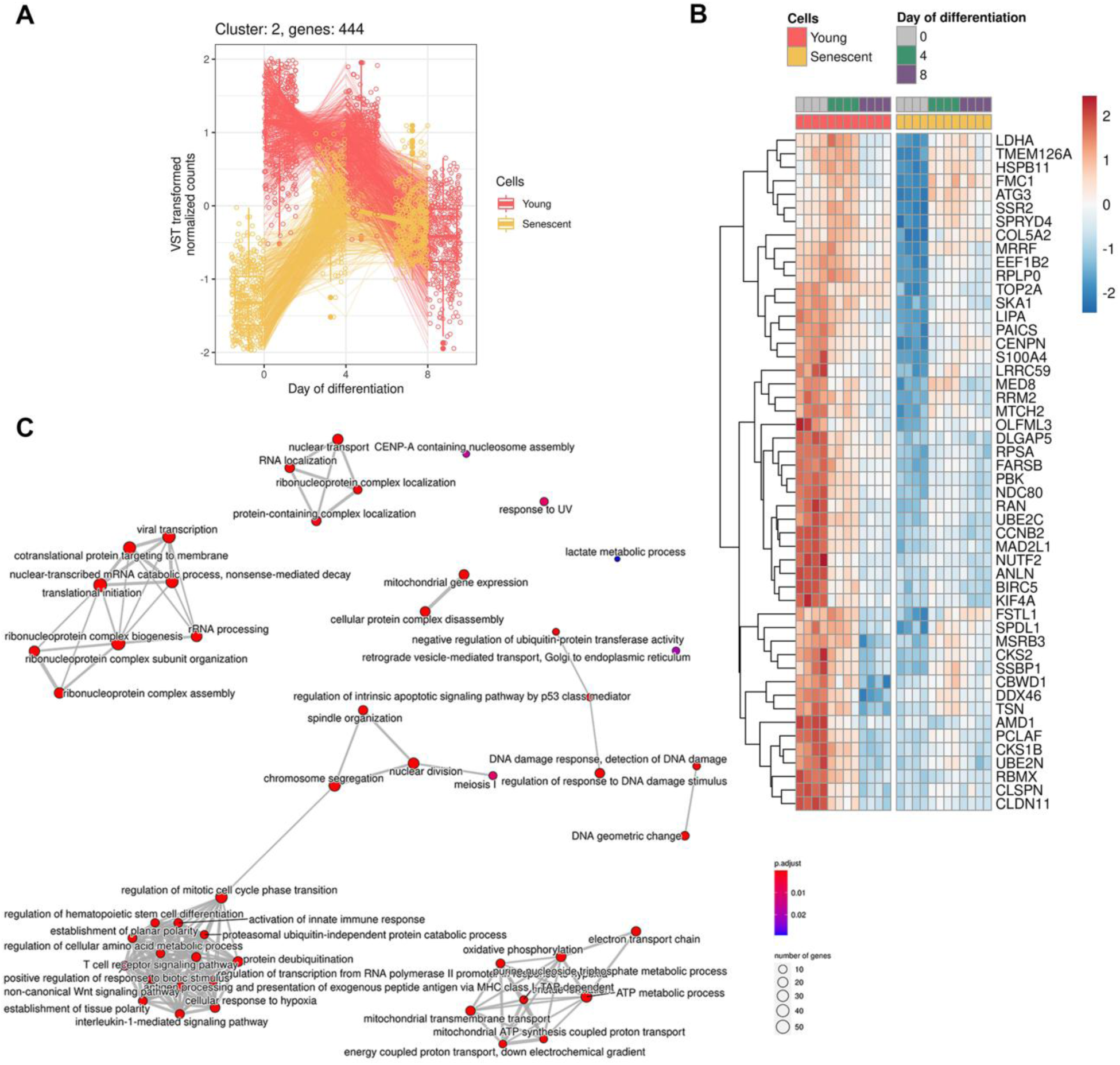
Cluster 2. A Expression of genes forming cluster B Heatmap reflecting expression of the top 50 DEGs in cluster C Functional enrichment analysis (FEA) of clustered genes in GO:BP terms

**Appendix Fig S3.**
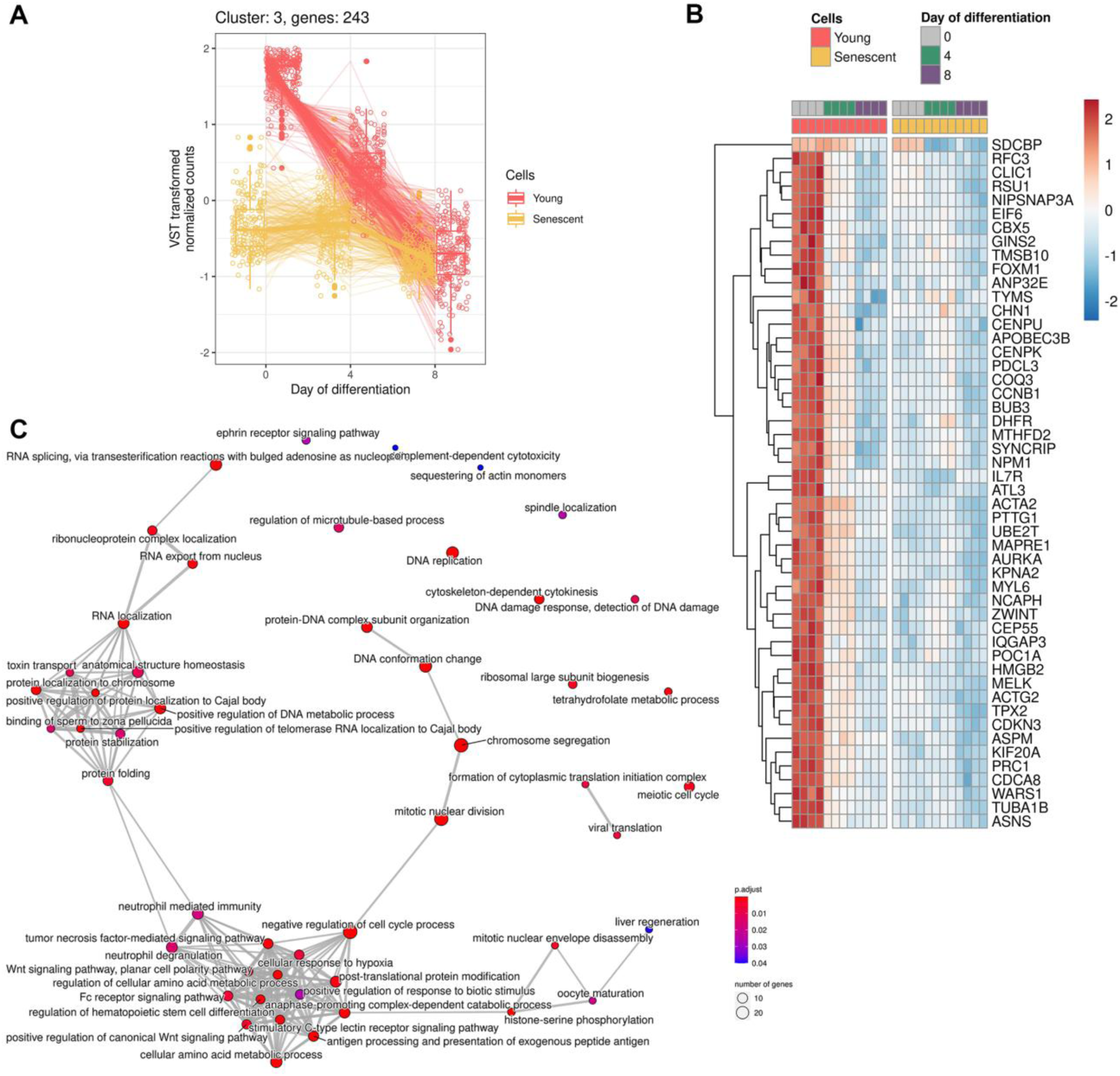
Cluster 3. A Expression of genes forming cluster B Heatmap reflecting expression of the top 50 DEGs in cluster C Functional enrichment analysis (FEA) of clustered genes in GO:BP terms

**Appendix Fig S4.**
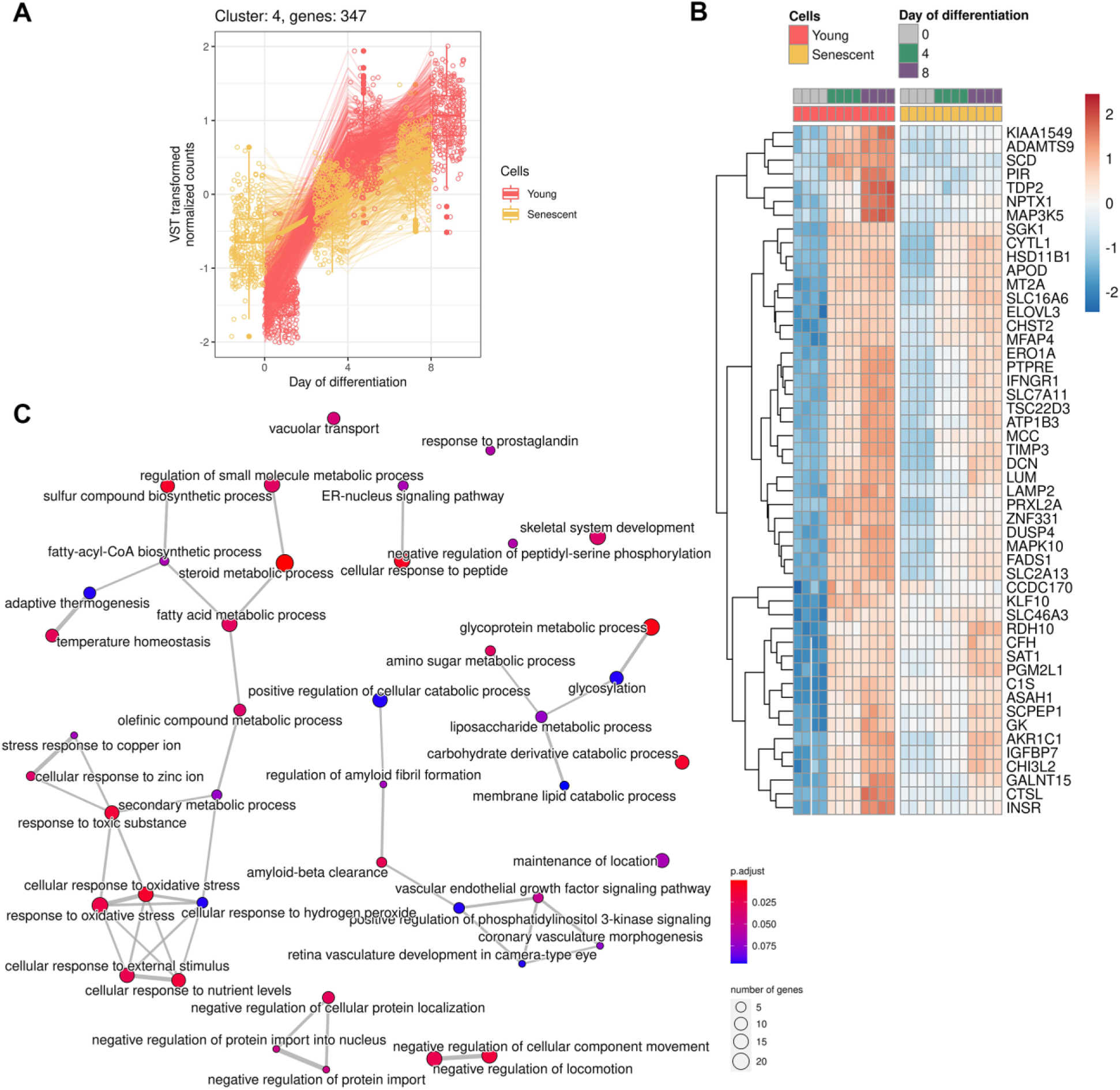
Cluster 4. A Expression of genes forming cluster B Heatmap reflecting expression of the top 50 DEGs in cluster C Functional enrichment analysis (FEA) of clustered genes in GO:BP terms

**Appendix Fig S5.**
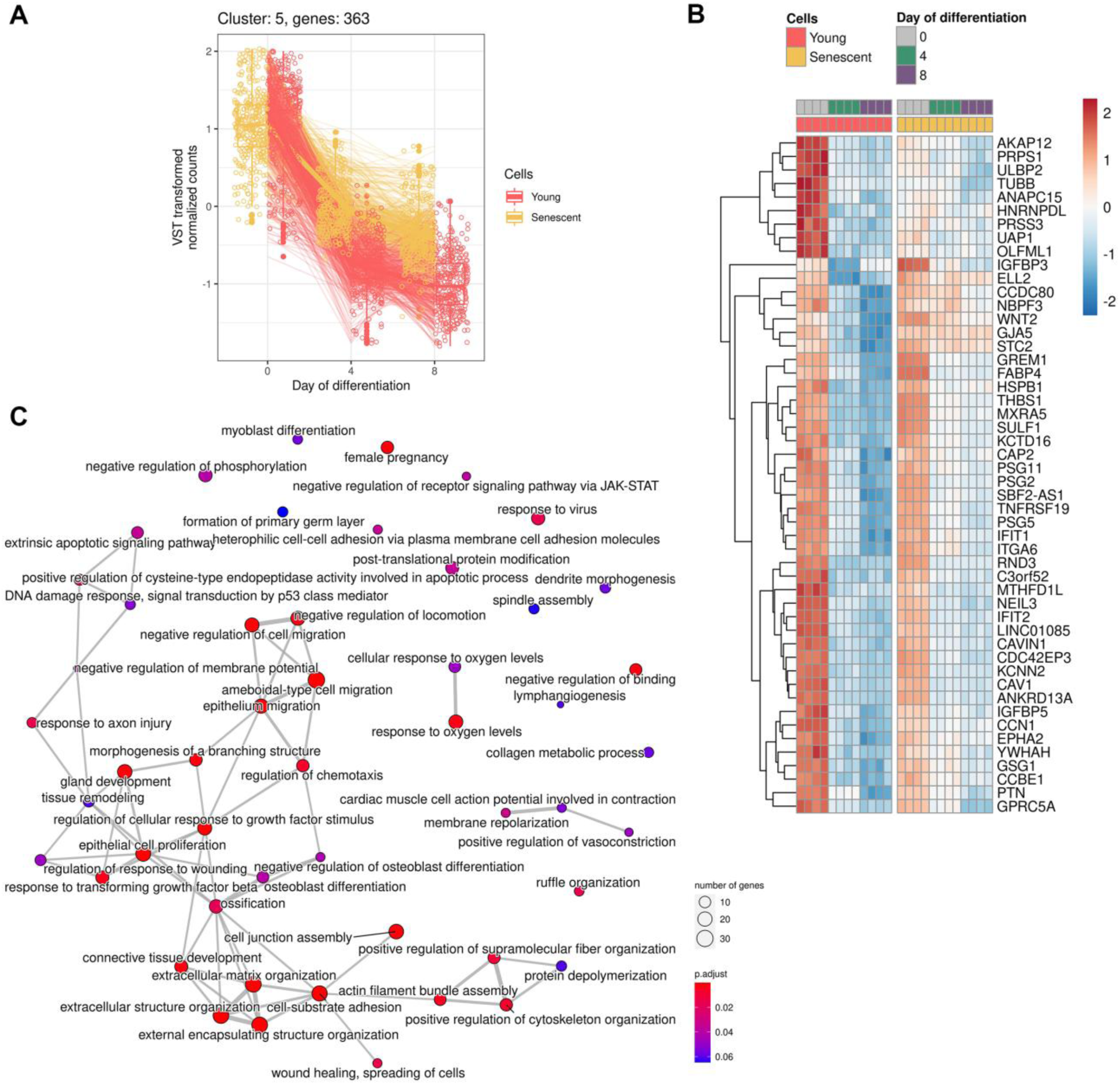
Cluster 5. A Expression of genes forming cluster B Heatmap reflecting expression of the top 50 DEGs in cluster C Functional enrichment analysis (FEA) of clustered genes in GO:BP terms

**Appendix Fig S6.**
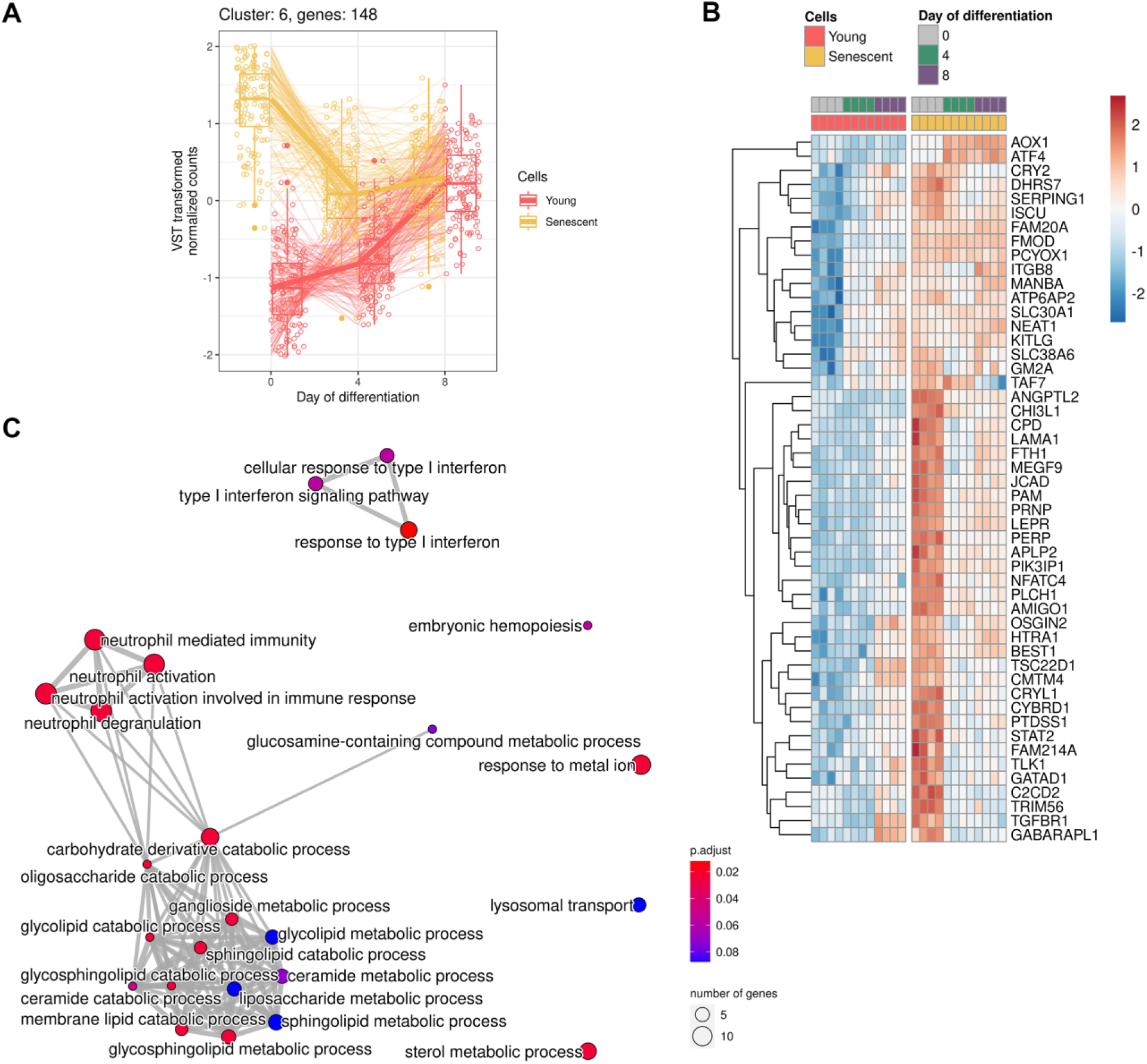
Cluster 6. A Expression of genes forming cluster B Heatmap reflecting expression of the top 50 DEGs in cluster C Functional enrichment analysis (FEA) of clustered genes in GO:BP terms

**Appendix Fig S7.**
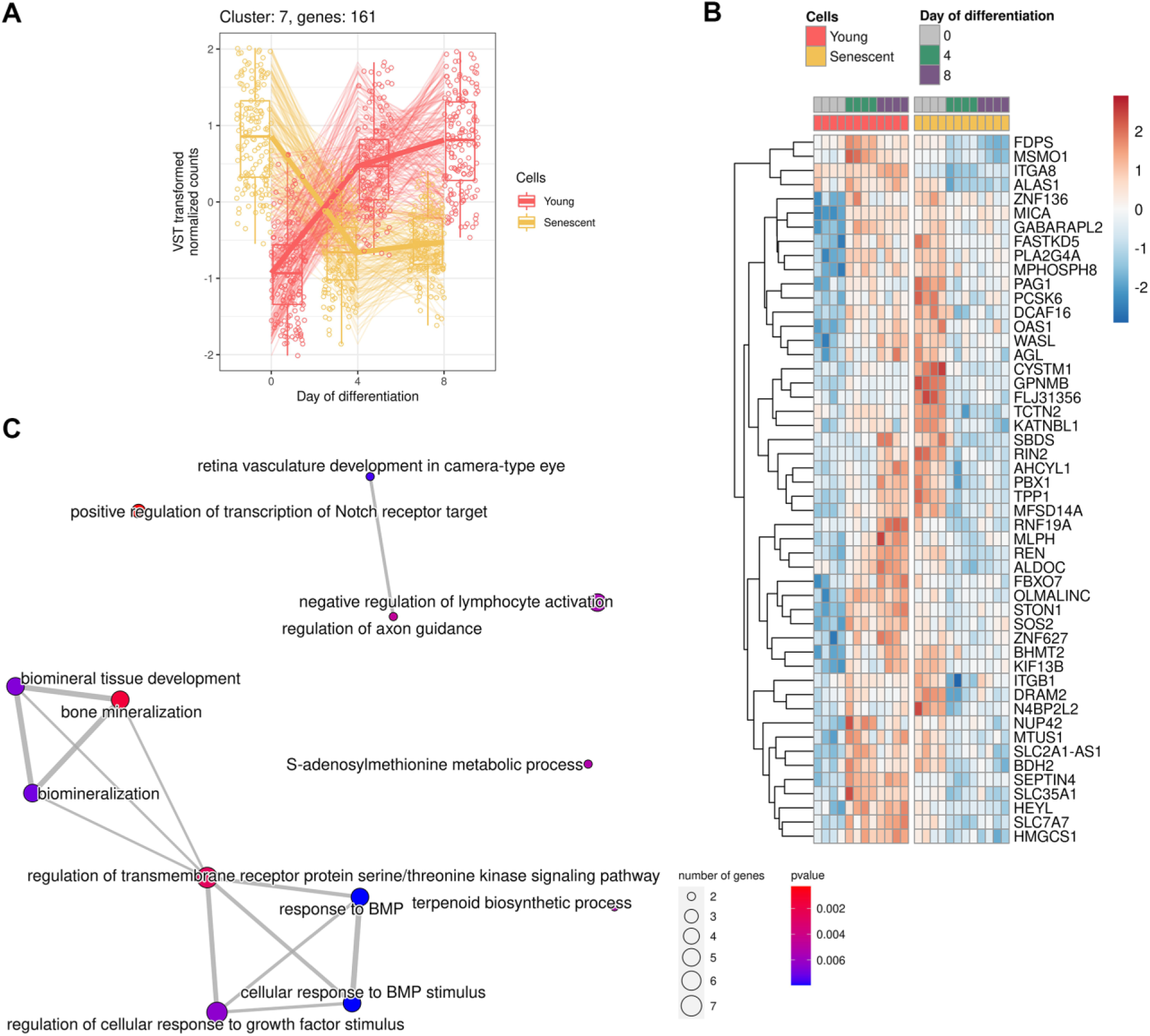
Cluster 7. A Expression of genes forming cluster B Heatmap reflecting expression of the top 50 DEGs in cluster C Functional enrichment analysis (FEA) of clustered genes in GO:BP terms

**Appendix Fig S8.**
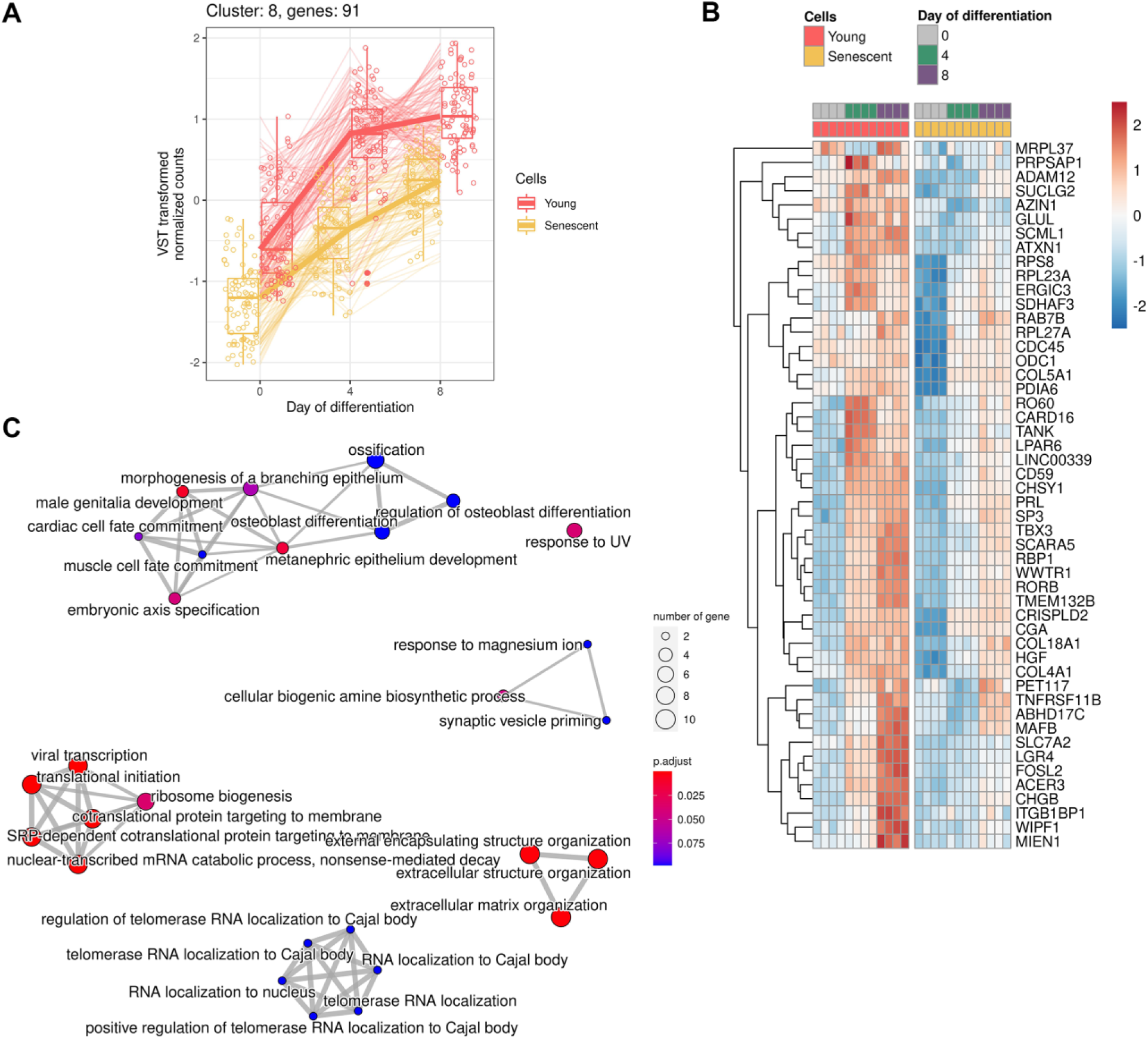
Cluster 8. A Expression of genes forming cluster B Heatmap reflecting expression of the top 50 DEGs in cluster C Functional enrichment analysis (FEA) of clustered genes in GO:BP terms

**Appendix Fig S9.**
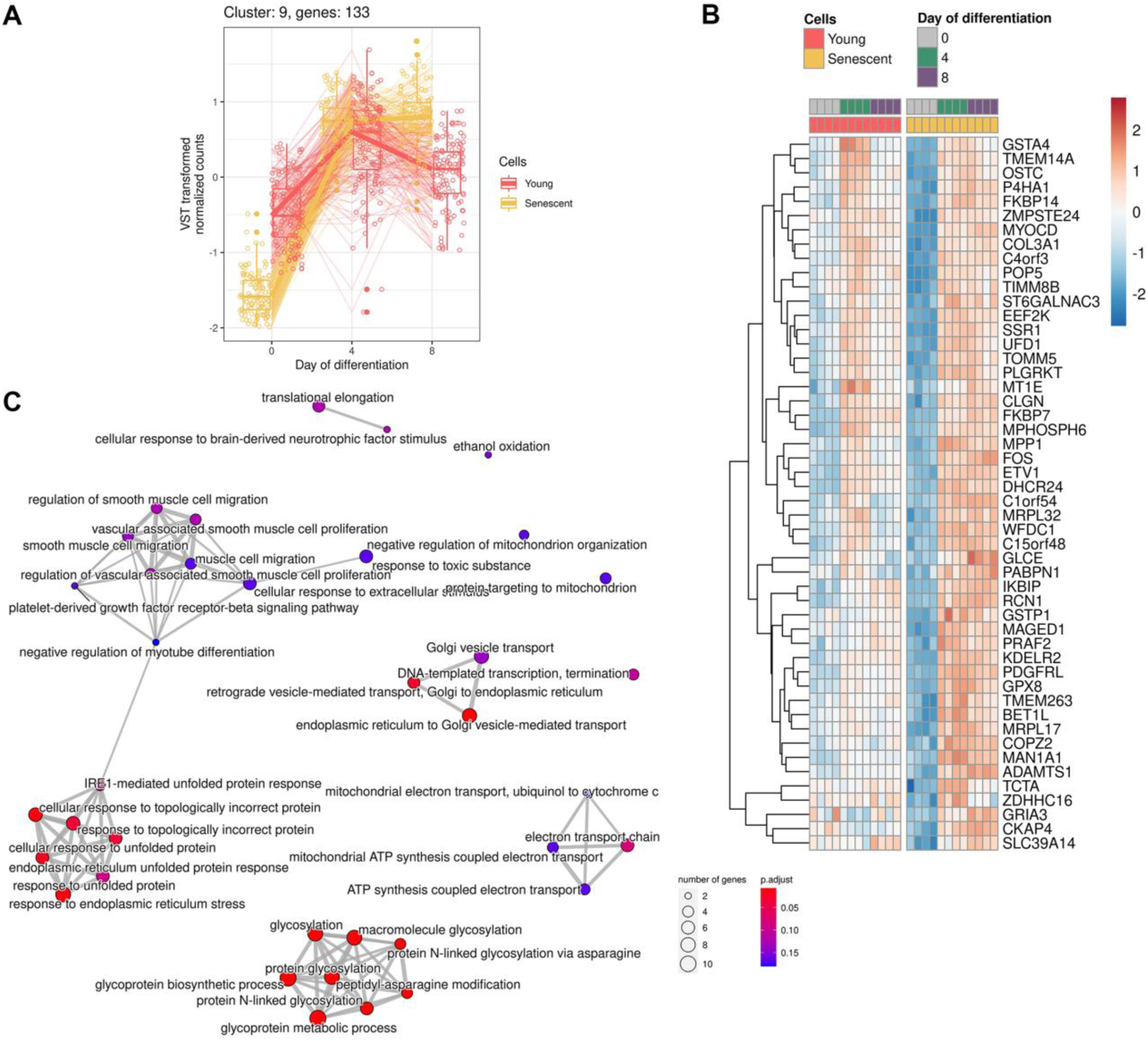
Cluster 9. A Expression of genes forming cluster B Heatmap reflecting expression of the top 50 DEGs in cluster C Functional enrichment analysis (FEA) of clustered genes in GO:BP terms

**Appendix Fig S10.**
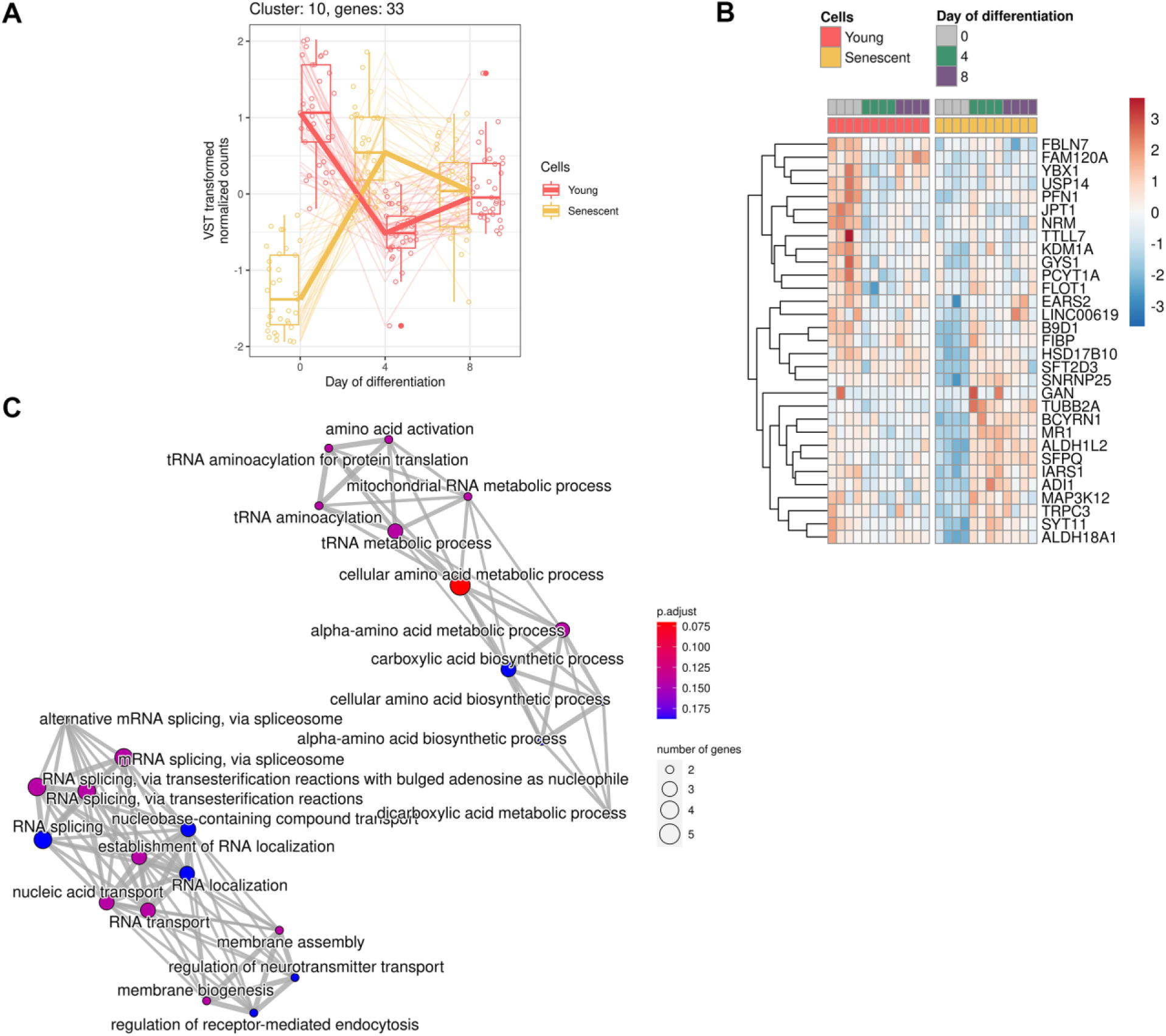
Cluster 10. A Expression of genes forming cluster B Heatmap reflecting expression of the top 50 DEGs in cluster C Functional enrichment analysis (FEA) of clustered genes in GO:BP terms

**Appendix Fig S11.**
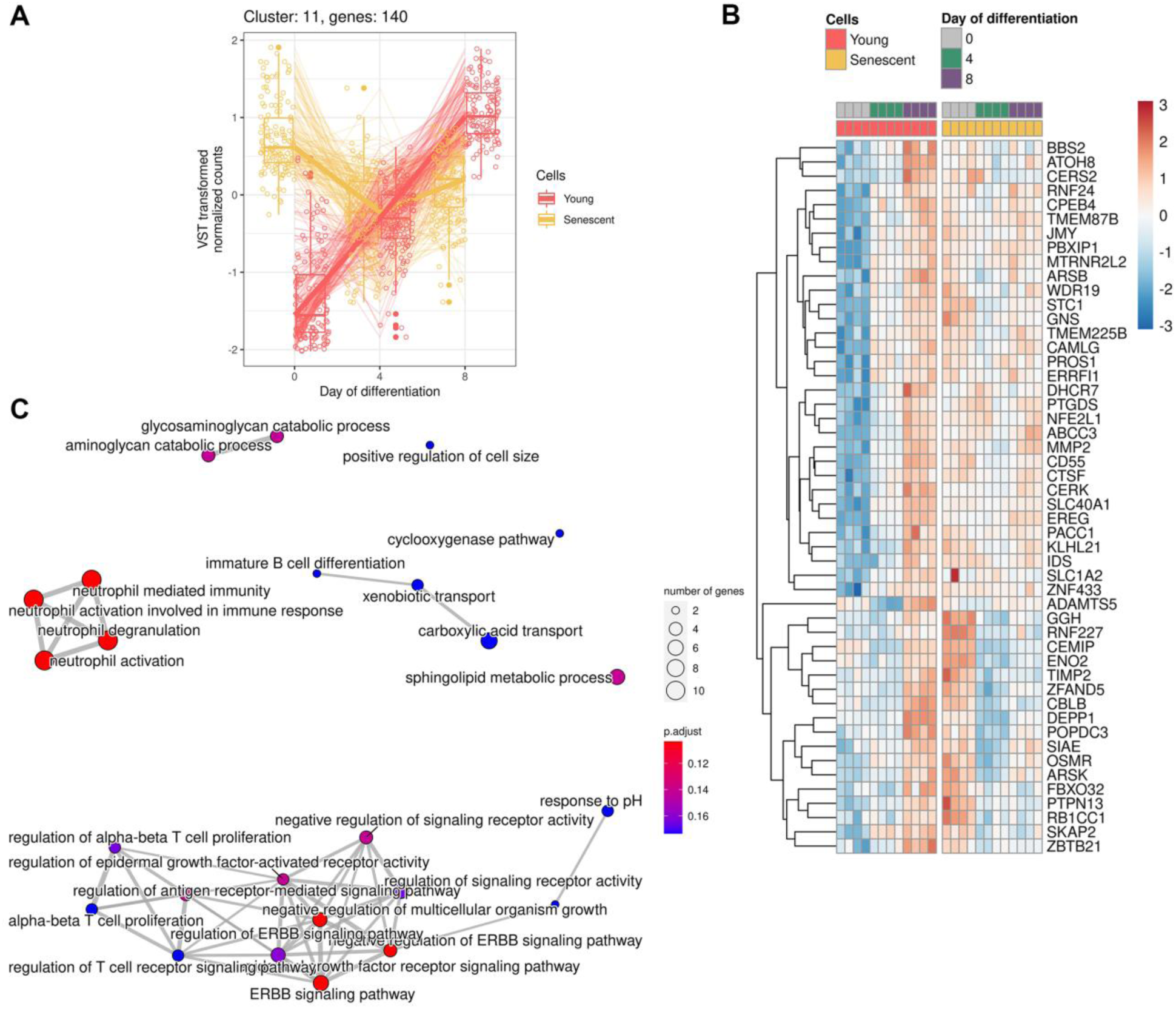
Cluster 11. A Expression of genes forming cluster B Heatmap reflecting expression of the top 50 DEGs in cluster C Functional enrichment analysis (FEA) of clustered genes in GO:BP terms

**Appendix Fig S12.**
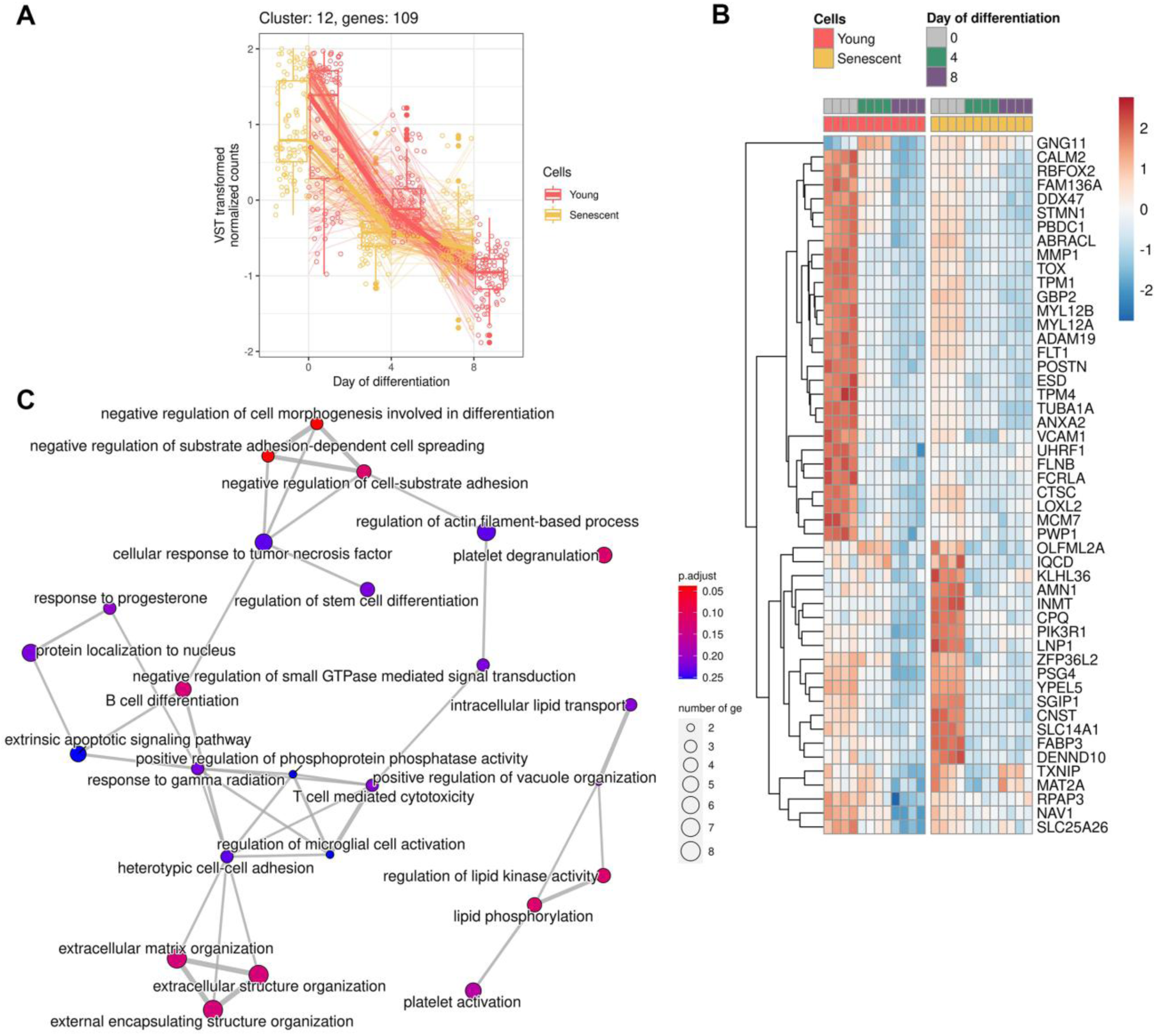
Cluster 12. A Expression of genes forming cluster B Heatmap reflecting expression of the top 50 DEGs in cluster C Functional enrichment analysis (FEA) of clustered genes in GO:BP terms

**Appendix Fig S13.**
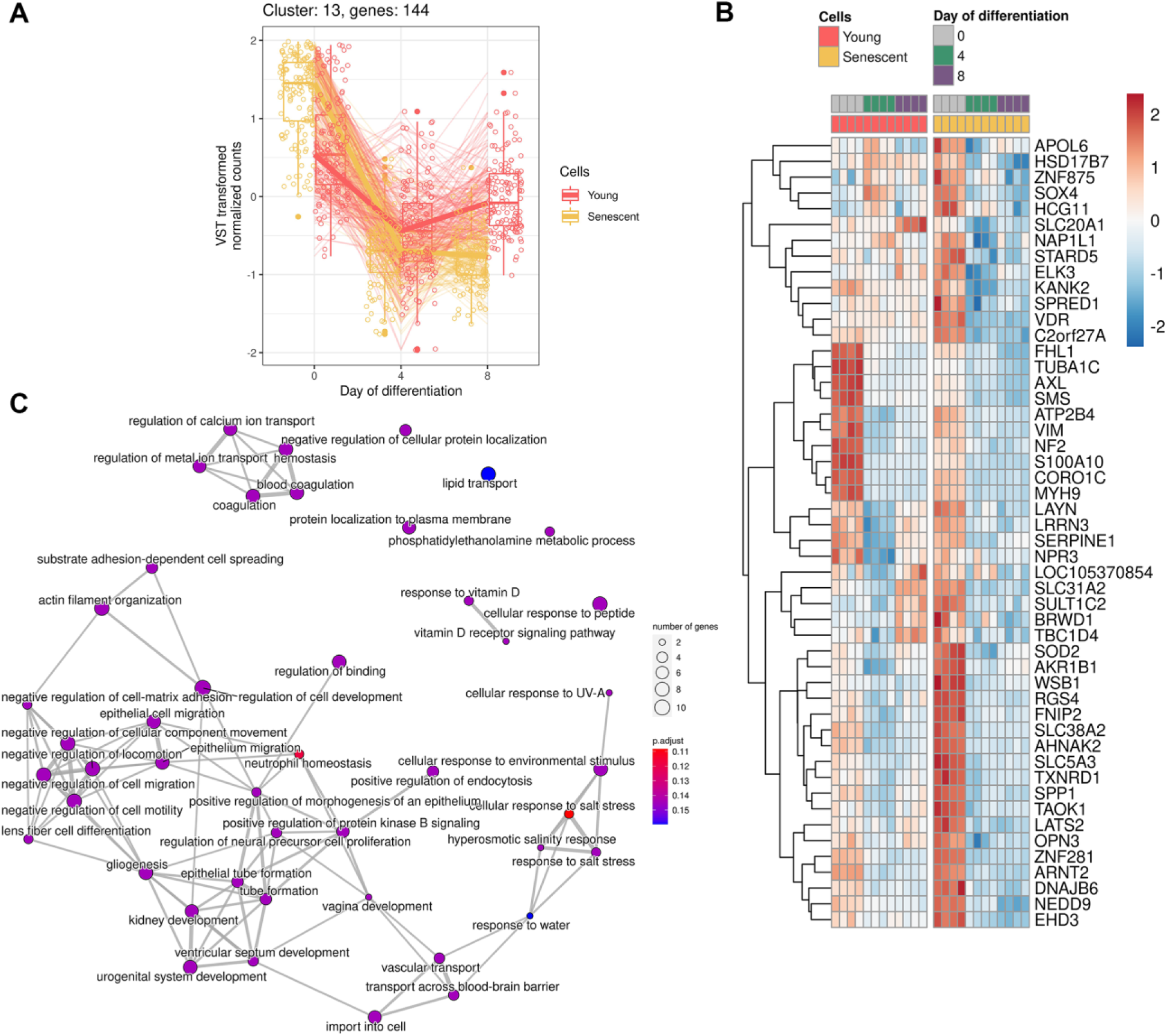
Cluster 13. A Expression of genes forming cluster B Heatmap reflecting expression of the top 50 DEGs in cluster C Functional enrichment analysis (FEA) of clustered genes in GO:BP terms

**Appendix Fig S14.**
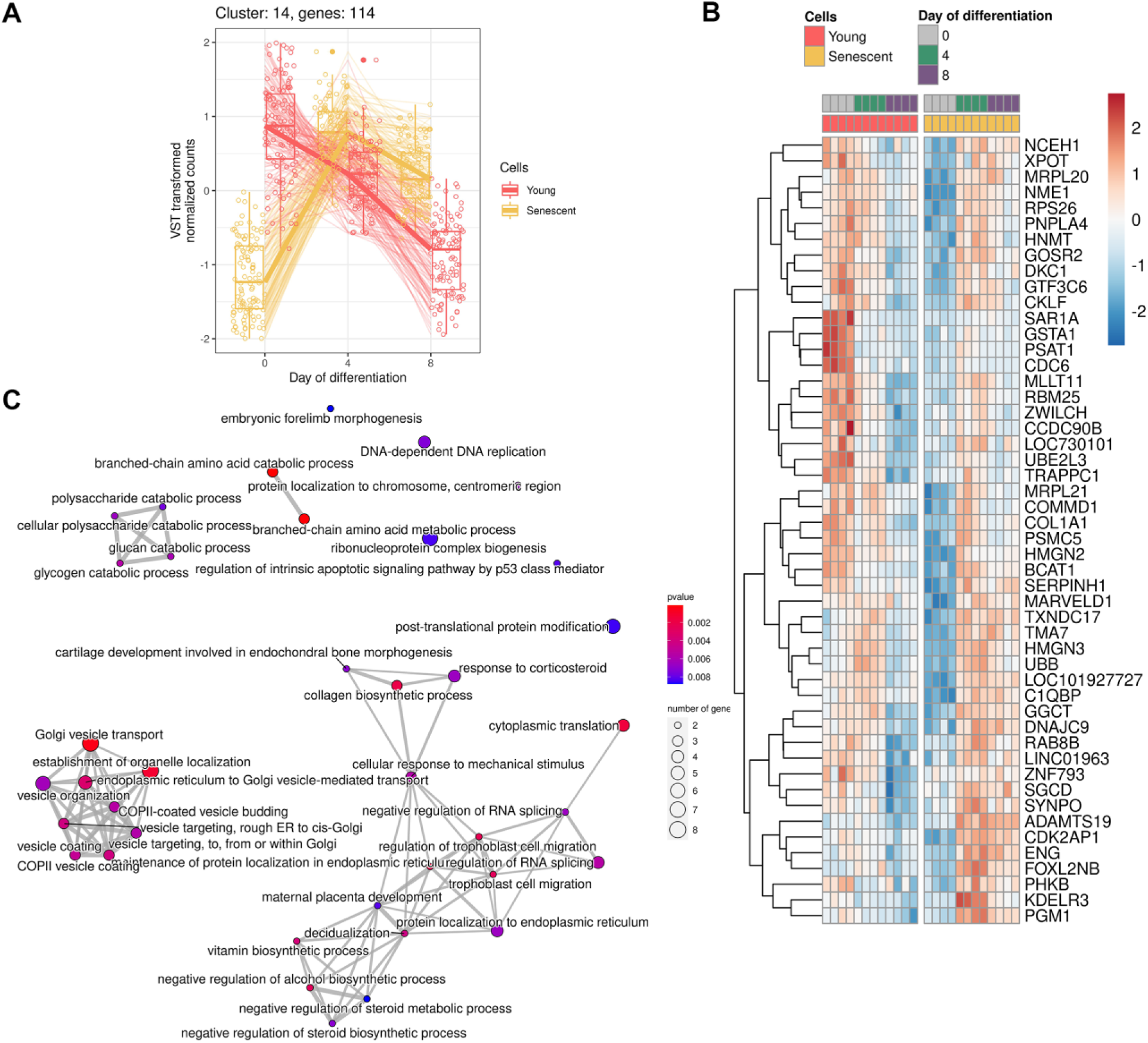
Cluster 14. A Expression of genes forming cluster B Heatmap reflecting expression of the top 50 DEGs in cluster C Functional enrichment analysis (FEA) of clustered genes in GO:BP terms

**Appendix Fig S15.**
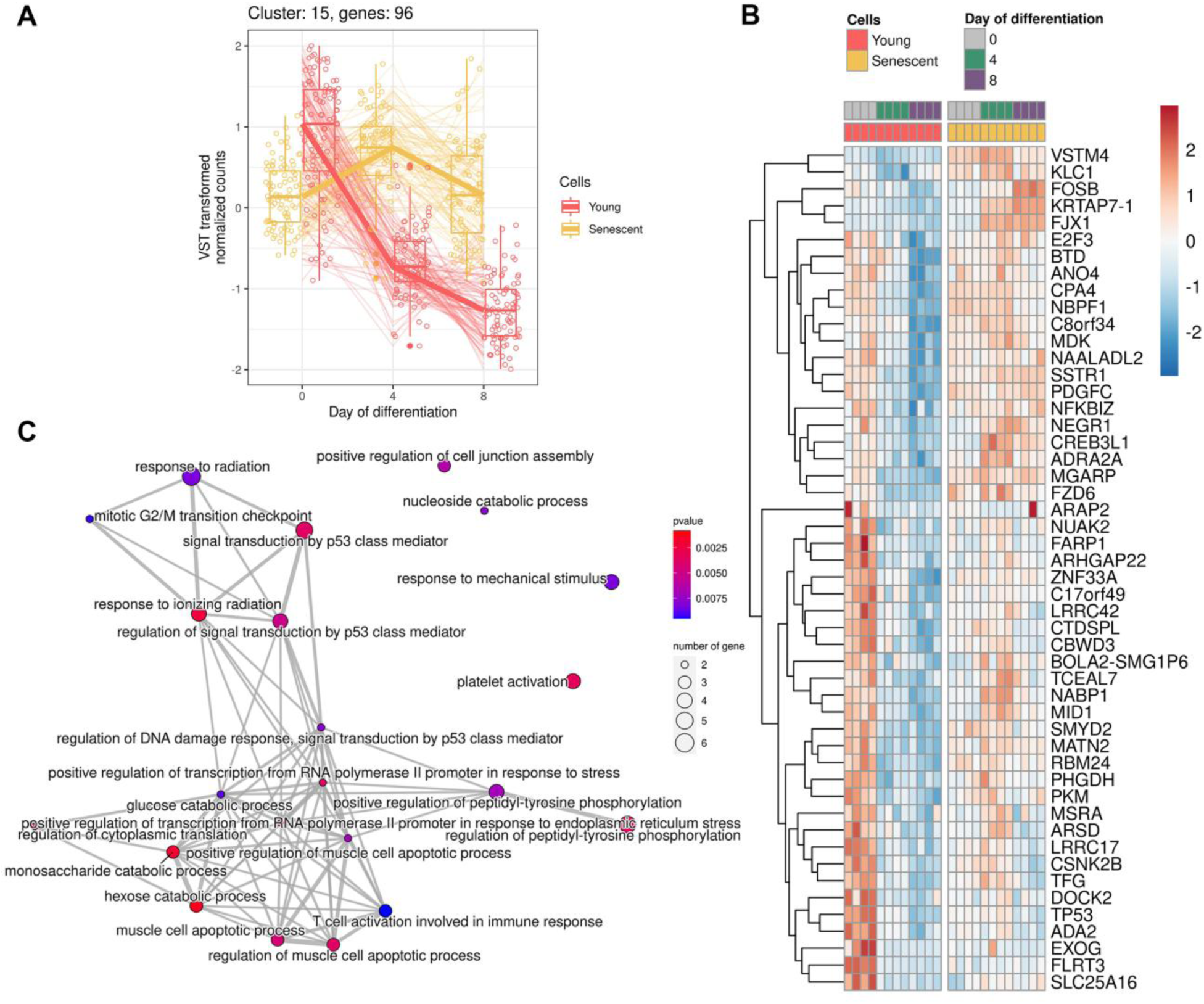
Cluster 15. A Expression of genes forming cluster B Heatmap reflecting expression of the top 50 DEGs in cluster C Functional enrichment analysis (FEA) of clustered genes in GO:BP terms

**Appendix Fig S16.**
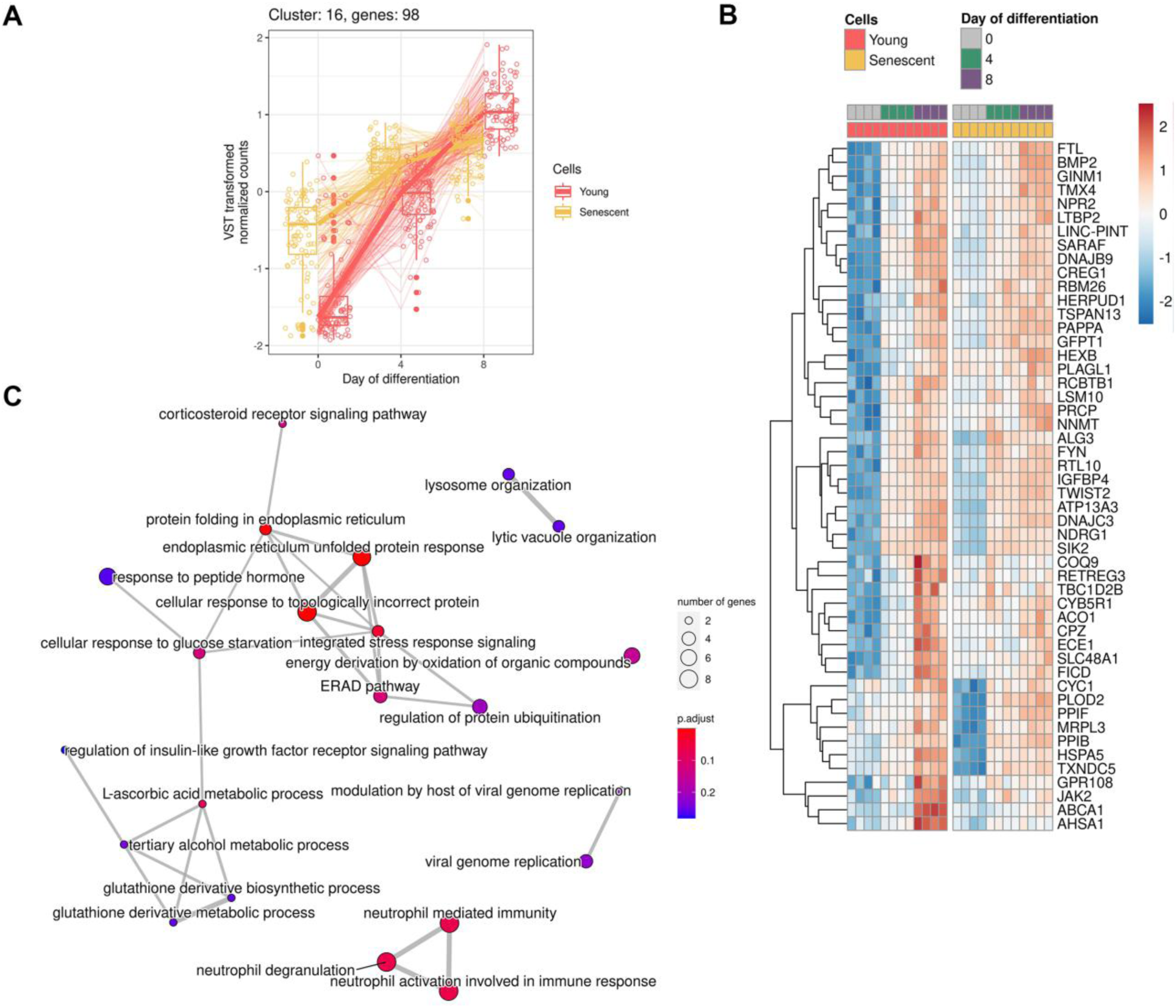
Cluster 16. A Expression of genes forming cluster B Heatmap reflecting expression of the top 50 DEGs in cluster C Functional enrichment analysis (FEA) of clustered genes in GO:BP terms

**Appendix Fig S17.**
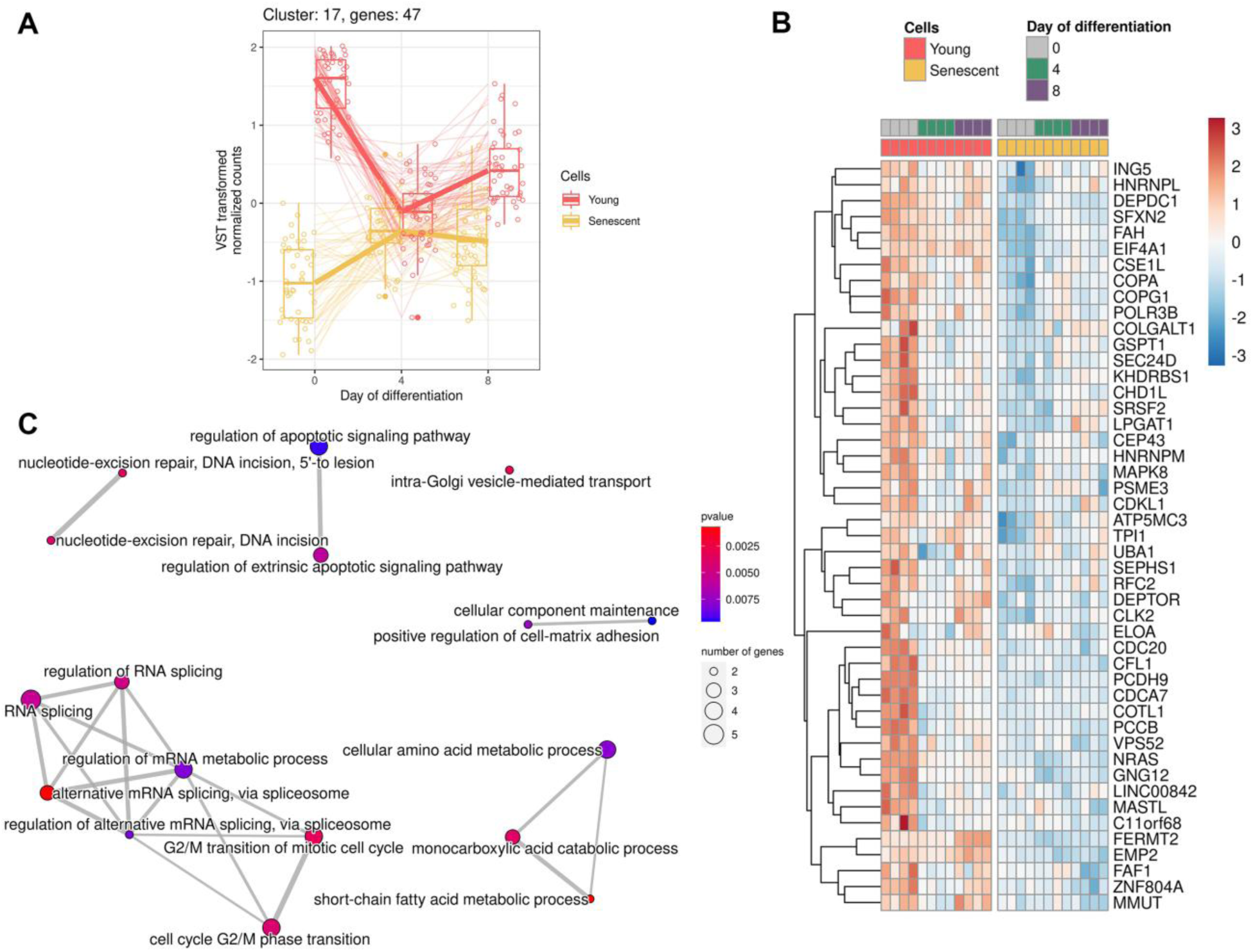
Cluster 17. A Expression of genes forming cluster B Heatmap reflecting expression of the top 50 DEGs in cluster C Functional enrichment analysis (FEA) of clustered genes in GO:BP terms

**Appendix Fig S18.**
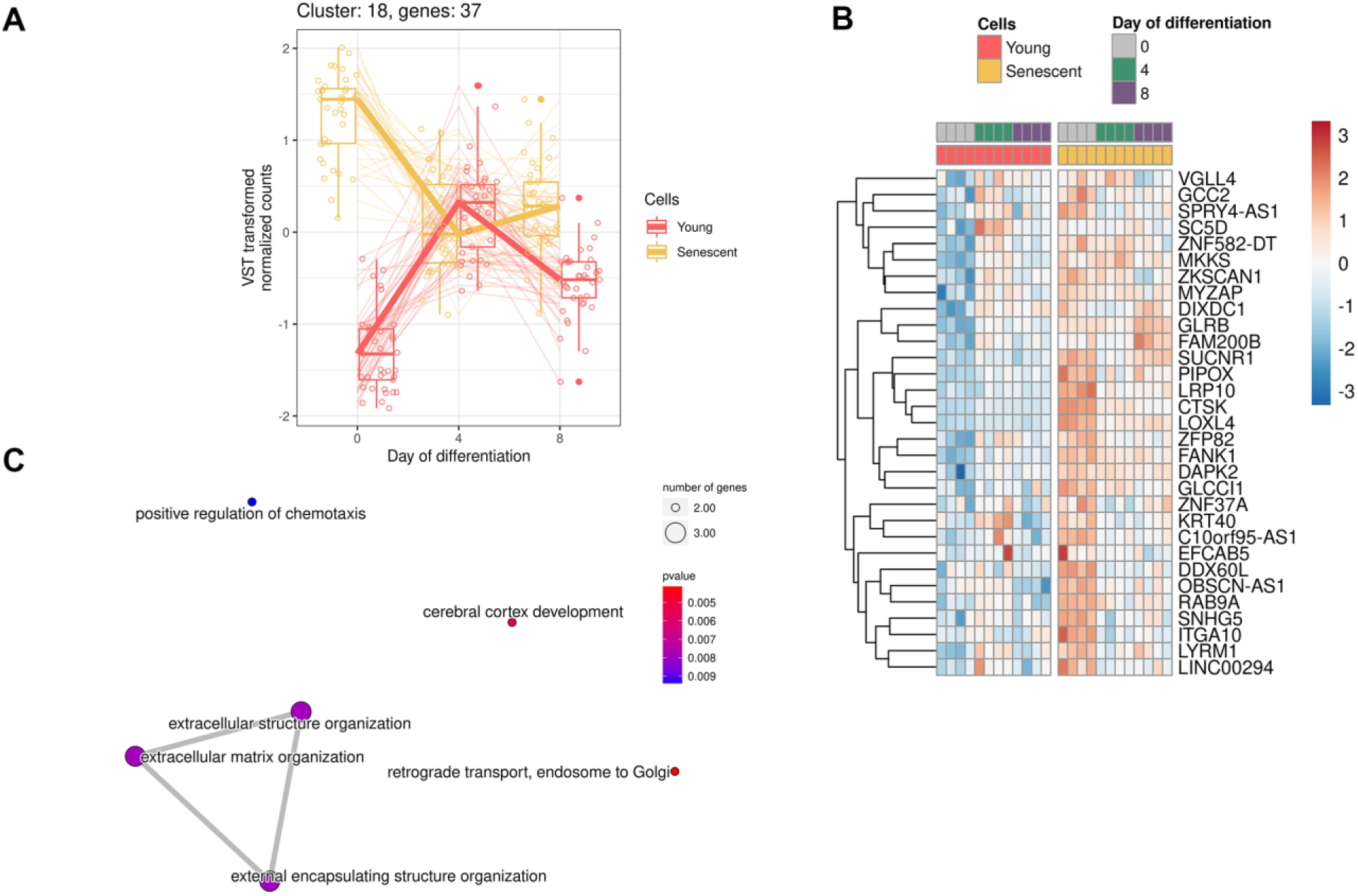
Cluster 18. A Expression of genes forming cluster B Heatmap reflecting expression of the top 50 DEGs in cluster C Functional enrichment analysis (FEA) of clustered genes in GO:BP terms

**Appendix Fig S19.**
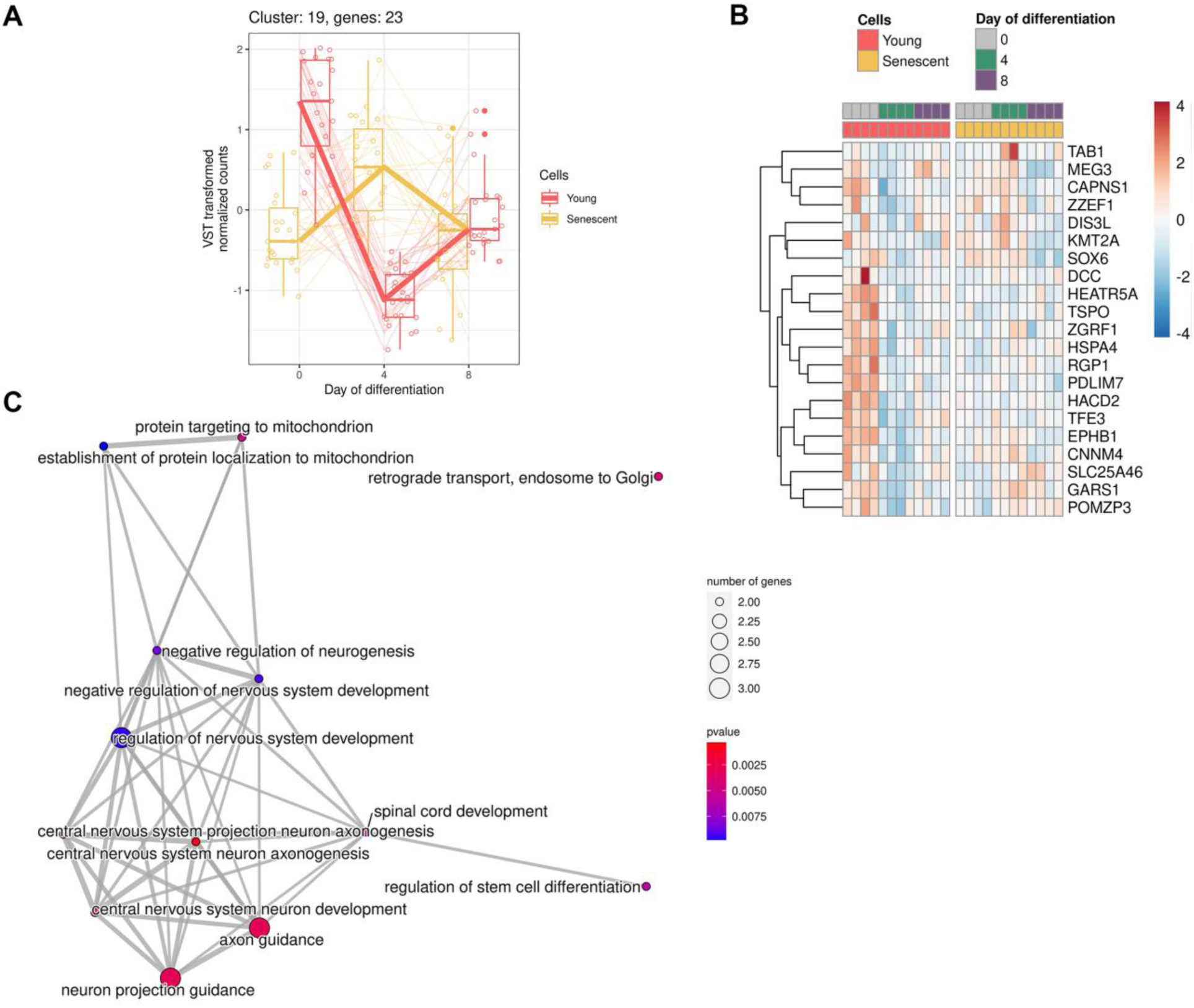
Cluster 19. A Expression of genes forming cluster B Heatmap reflecting expression of the top 50 DEGs in cluster C Functional enrichment analysis (FEA) of clustered genes in GO:BP terms

**Appendix Fig S20.**
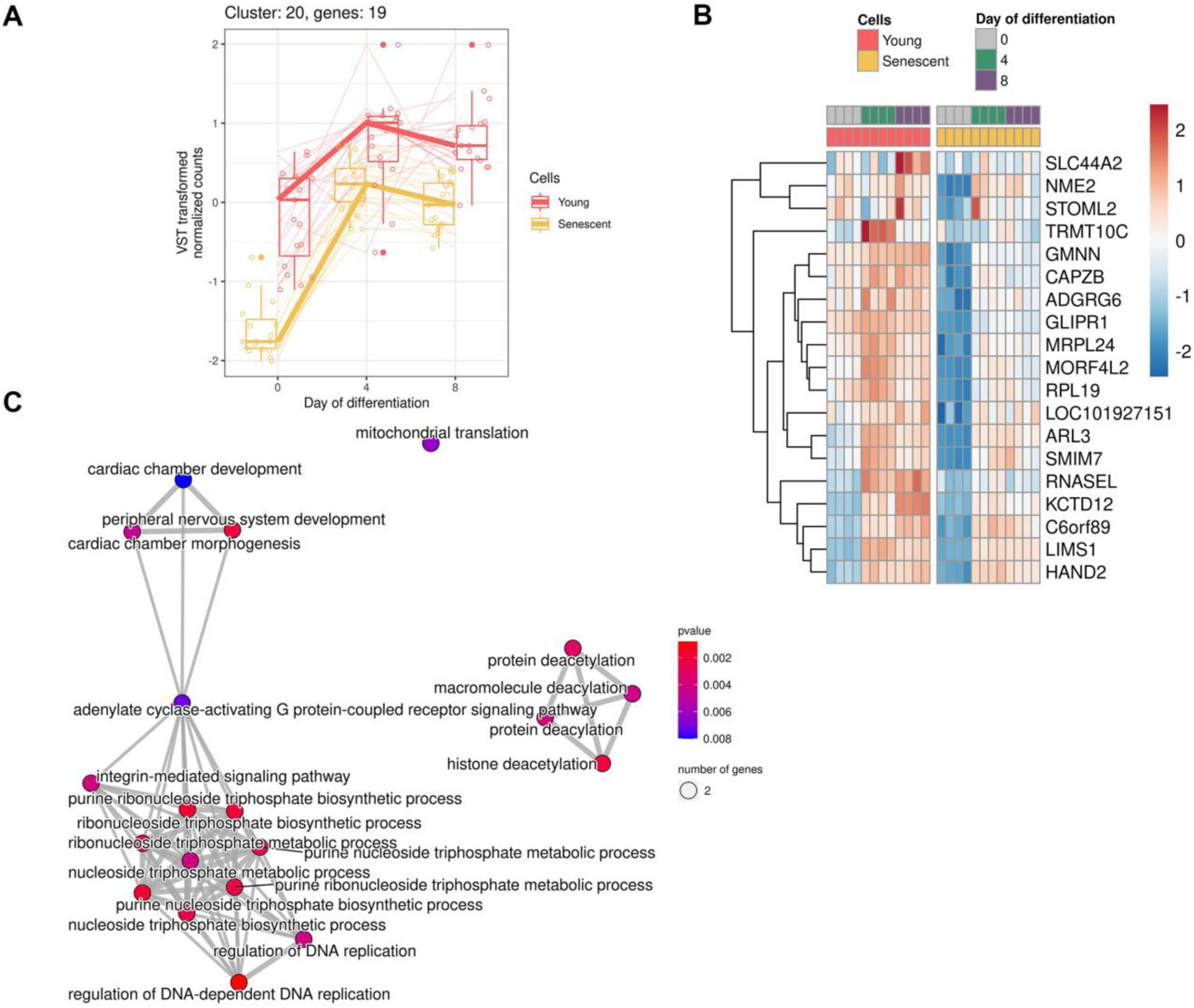
Cluster 20. A Expression of genes forming cluster B Heatmap reflecting expression of the top 50 DEGs in cluster C Functional enrichment analysis (FEA) of clustered genes in GO:BP terms

**Appendix Fig S21.**
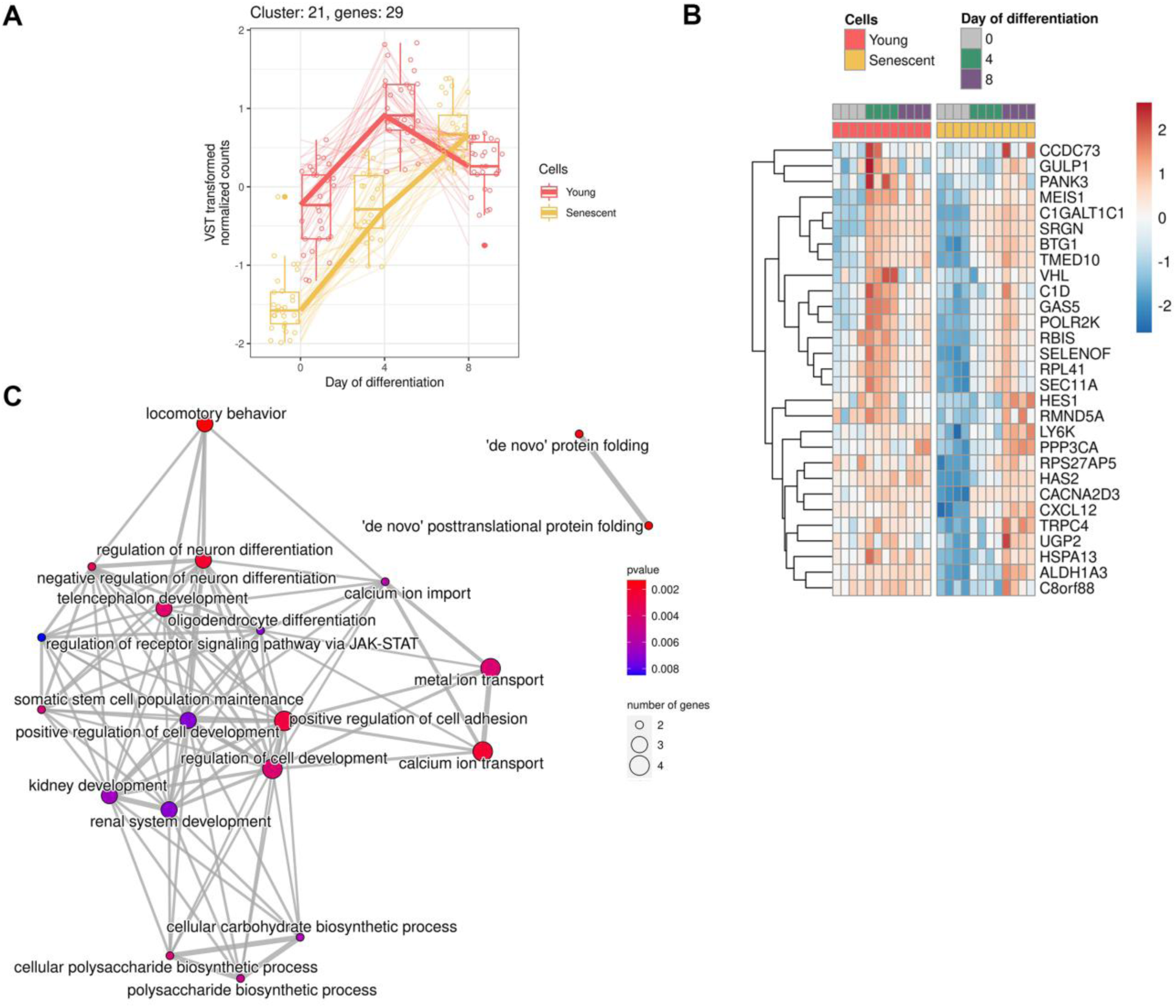
Cluster 21. A Expression of genes forming cluster B Heatmap reflecting expression of the top 50 DEGs in cluster C Functional enrichment analysis (FEA) of clustered genes in GO:BP terms

**Appendix Fig S22.**
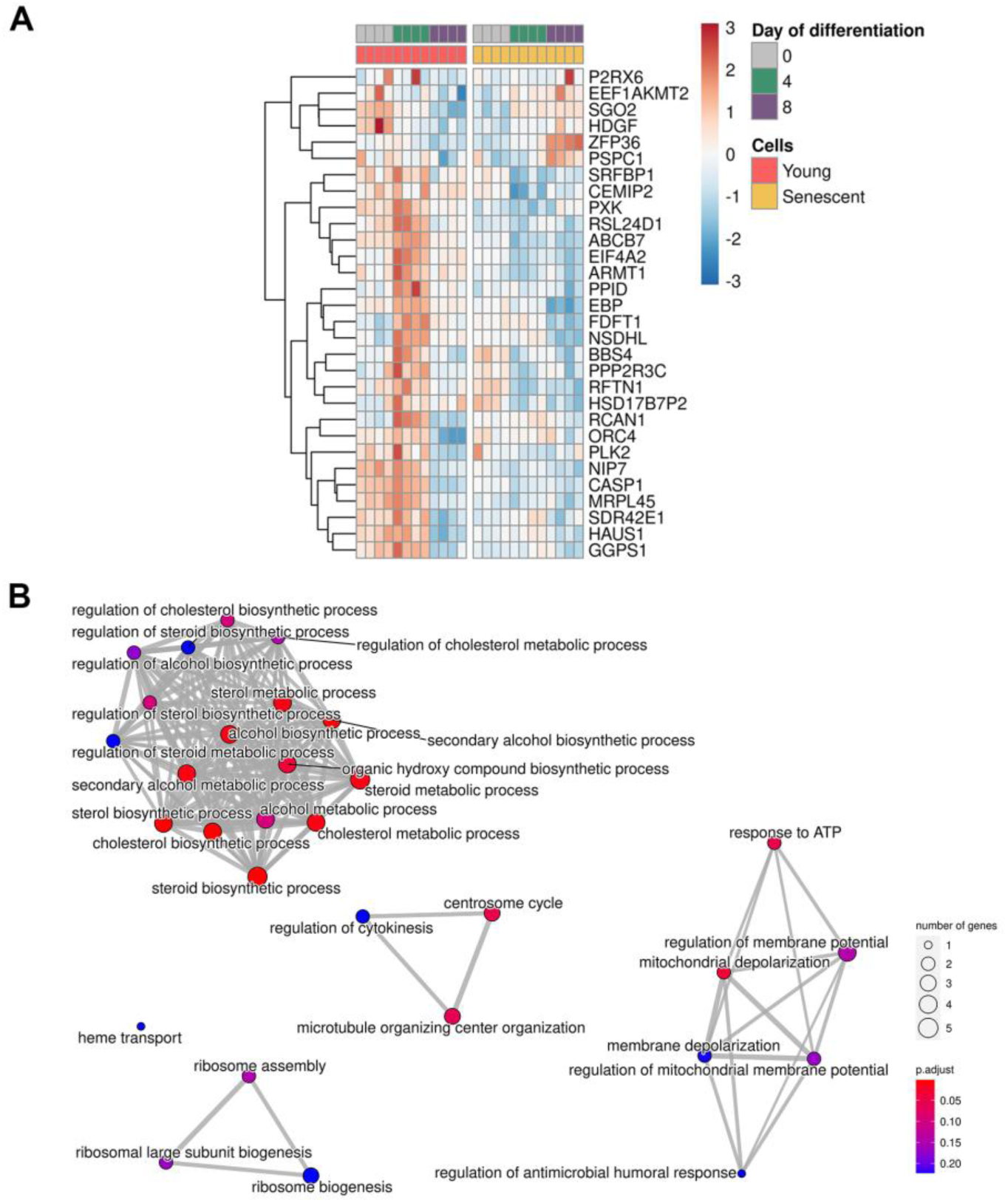
Non-clustered genes. A Expression of genes forming cluster B Heatmap reflecting expression of the top 50 DEGs in cluster C Functional enrichment analysis (FEA) of clustered genes in GO:BP terms

